# Rare variants drive high variance in human ancestral fitness at mutation-selection-drift balance

**DOI:** 10.64898/2026.07.29.741368

**Authors:** Ulises Hernández, Walid Mawass, Joseph Matheson, Joanna Masel

## Abstract

It is an open question whether variation in the genetic load of unconditionally deleterious mutations contributes substantially to the variability in human disease. Here, we solve for mutation-selection-drift balance and predict variation in genetic load given a realistic human genome-wide deleterious mutation rate, *U*, and a distribution of fitness effects (DFE). Empirical estimates of *U* come from sequence constraint, which fails to count slightly deleterious mutations that nevertheless fix. We use the inferred DFE to correct for this and conclude that total human *U*>3.8. Two humans typically differ in ancestral fitness by 17-33% given uncertainty in *U*, or by 6-49% when we consider a broad range of alternative DFEs. Results are similar for other species with larger mean selection coefficients, such as other mammals. Most variation in load comes from rare variants with frequencies below 1%, with a substantial fraction coming from ultra-rare variants below 0.01%. This could help explain why some of the heritability observed in pedigree studies is missing from genome-wide association studies. Accounting for rare and ultra-rare variants, e.g., via variant-effect prediction of unique mutations from whole-genome sequencing rather than via polygenic risk scores, could help identify individuals at high risk of disease.

**Significance:** Many human mutations mildly disrupt molecular function, e.g., by destabilizing proteins. Having too many of these mutations would have reduced fitness in ancestral human environments and might contribute to disease today. Here, we mathematically derive how much variation in fitness such mutations cause, using estimated human parameter values. Rare variants with larger fitness effects contribute the most. Identifying individuals with high disease risk likely requires methods capable of scoring rare variants.

## Introduction

Genetic variants can predispose individuals to disease (1, 2). Genetic screening to identify high-risk individuals can be helpful for diseases that benefit from earlier intervention (3, 4). The usual approach is to construct a polygenic risk score (PRS) based on common variants identified through genome-wide association studies (GWAS) (3, 5). Emerging approaches incorporate rare variants into PRS by performing GWAS on whole-genome sequences (6) or by using sequence conservation among primates to predict variants of larger effect, even when they have never been observed before (7, 8).

Genetic variants that promote disease can be due to a mismatch between past environments and the current environment (9, 10), to genetic trade-offs where mutations increase fitness at the expense of increasing disease risk (9), or to deleterious mutations that have not yet been purged. A variety of empirical approaches suggest widespread purifying selection on disease-associated genetic variants (11). Alleles with large effects within a GWAS tend to be at a lower frequency (12–15), to be younger (16), to be found in genomic windows with intensified background selection (16, 17), and to be found at more conserved sites (15, 17, 18).

Purifying selection can occur, for example, when mutations break molecular functionality, e.g., by destabilizing protein folding (19), making them broadly deleterious across most environments. Many such mutations segregate in populations, leading to a genetic “load” of deleterious mutations (20–23).

While some deleterious mutations have highly specific effects on a single disease, many are likely to have diffuse effects on whole-organism physiology. The aggregate deleterious load across all sites can thus be thought of as itself a trait, one that is likely to correlate with a broad range of diseases. A single mutation with a substantial deleterious selection coefficient might have many small effects that are each too small to be significant for a given disease, but summing effects over many such deleterious mutations might nevertheless predict that same disease.

The higher the variance in total deleterious load, the more plausible it is that it drives variation in disease. We therefore revisit calculations of the magnitude of variance in load, using both improved calculations and improved parameter value estimates.

Load is often expressed not as variance, but as a difference between two genotypes. Classical studies focused on “lag load” (20–22, 24, 25), which is the difference between the mean population fitness and that of an unrealistic genotype with zero deleterious mutations. This yields a “mutation load paradox” (26) of absurdly high lag load (23, 27–29). The non-existence of a mutation-free individual makes lag load a misleading concept (30, 31). A better reference genotype is that of the best individual actually present in the population (30, 32). Logically, the expected number of viable offspring of the best individual 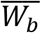 must be compatible with the species’ life history. For human life history, (33) argued that maximum fitness 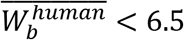.

Most previous models of load solved for mutation-selection balance (MSB). Variation in load then depends on two parameters: the whole-genome deleterious mutation rate, *U*, and the mean heterozygous selection coefficient of a new deleterious mutation, 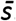. Under co-dominance, relative fitness at MSB is log-normally distributed with mean 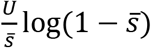 and variance 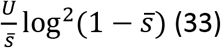.

We use 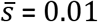 as an upper bound based on a distribution of fitness effects (DFE) inferred from the site frequency spectrum of non-synonymous mutations from the ESP human sample in (34). (The LuCamp sample in the same study provides a lower 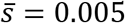 estimate). As a lower bound, we use 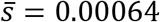, which corresponds to the mean of the DFE of ultra-conserved noncoding regions inferred by (35). Even our upper bound 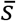 is small enough such that 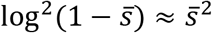, making the variance in log relative fitness 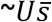 (Equation 10 in (36)).

*U* seems to be high in many species, especially humans (30, 37, 38). *U* can be estimated as *U*_*obs*_ = 2*μG*_*del*_*P*_*del*_ where 2*μ* is the per-diploid-site mutation rate, *G*_*del*_ is the genome-wide number of diploid sites subject to deleterious mutations, and *P*_*del*_ is the probability that a new mutation at one of these sites is deleterious (33, 39). (33) assumed neutral evolution of regions that are rich in transposable elements that are shared with chimpanzee and hence presumed inactive. These “ancestral repeat regions” make up 45% of the human genome (40). From the human diploid genome size of 3 × 10^9^, (33) calculated *G*_*del*_ = 3 × 10^9^ × (1 − 0.45) = 1.65 × 10^9^. (41) estimated *P*_*del*_ = 0.057 from the extent to which the rest of the genome evolves more slowly than ancestral repeat regions. Here, we use *μ* = 1.515 × 10^−8^ which combines point mutations at a rate 1.341 × 10^−8^ (42) with indels at a rate 0.174 × 10^−8^ (42). This yields 2.85, which we argue below is a lower bound, giving *U*_*obs*_ > 2.85.

To obtain an upper bound on *U*_*obs*_, we estimate 2*μG*_*cons*_ (43), where *G*_*cons*_ is the genome-wide number of diploid individual nucleotide sites subject to some degree of purifying selection. *G*_*cons*_ is estimated as the number of identifiable sites that evolve at a significantly lower rate than ancestral repeats. Using the whole-genome alignment of 240 mammal species, (44) estimated *G*_*cons*_ = 10.7%. Treating all mutations in *G*_*cons*_ as deleterious yields *U*_*obs*_ < 9.73. While this estimate is deflated by ignoring deleterious mutations outside *G*_*cons*_, we nevertheless consider it an upper bound because it assumes that all mutations in constrained regions are strongly deleterious.

Combining *U*_*obs*_ = [2.85 − 9.73] with 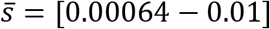 implies that under MSB, load makes the 75th-percentile fittest individual 6-52% fitter in an ancestral human environment than a 25th-percentile individual. Variance this high suggests that load is an important component of human trait variation.

Drift has two opposing effects on load variance. First, drift might increase the frequency of segregating deleterious alleles above the MSB expectation. This increases the variance in fitness, as has previously been studied in the context of increasing lag load (24, 25). Second, drift increases allele loss, which reduces the variance in fitness. Previous models ignored the latter effect by using diffusion theory, which does not assign a probability for a site to be monomorphic. This is problematic for human genomes, where most sites are expected to be monomorphic because new mutations arrive only every 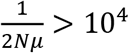 generations (using *μ* as estimated above, and ancestral *N* ≈ 10,000). Here we account for monomorphic sites under mutation-selection-drift balance.

With drift, the entire DFE matters, not just 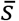. Our main reference point is the non-synonymous DFE from the ESP human sample in (34), a gamma distribution with mean 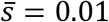 and coefficient of variance CV=2.43. (The LuCamp sample in the same study has mean 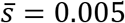 and CV=2.15.) At the other extreme, an estimated DFE of conserved non-coding regions has 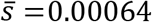 and CV=2.92. We consider a range for 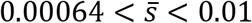. Given greater uncertainties in non-coding inference, we use the higher missense value, CV=2.43, by default. A mixture of site classes is likely to increase the CV. For consistency with the ESP DFE inference, we model drift with *N* = 11,300.

We also revisit estimates of human *U. U*_*obs*_ underestimates *U* for four reasons (45, 46). First, beneficial mutations are ignored, thereby deflating the estimation of *P*_*del*_ from constraint. Second, deleterious mutations outside *G*_*del*_ are ignored. Third, the divergence estimates underlying *P*_*del*_ were not free of polymorphisms, deflating *P*_*del*_.

Fourth, which we address here, is that *P*_*del*_ = 0.057 ignores weakly deleterious mutations. Our use of *P*_*del*_ to derive *U*_*obs*_ follows the neutral theory of molecular evolution, which assumes that deleterious mutations never fix. Because inference of the DFE points to many weakly deleterious mutations (34, 35), this procedure will underestimate *U*.

Here, we first revise the estimation of human *U*, focusing on this fourth challenge. We also replace models that balance irreversible mutation and selection (36, 47) with models of reversible mutation, selection, and drift. We then combine estimates of *U* and the DFE within our new model to derive the magnitude of fitness variance (in the ancestral human environments that shaped the site frequency spectrum from which the DFE is inferred) attributable to unconditionally deleterious mutations.

## Model

### *U* estimation

The DFE of deleterious mutations includes many mutations that are too weakly deleterious for selection to reliably prevent their fixation. To the degree that they fix at neutral rates, they do not contribute to our *U*_*obs*_ estimate. *U*_*obs*_ captures only those point mutations absent from divergence data because they were deleterious. That is,

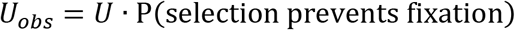

where P() refers to probability. We quantify this probability by comparing to a counterfactual with no selection:

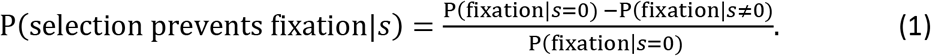

For a new mutation in a diploid population of constant size *N*, 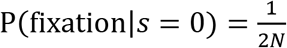. We model selection by allowing each selected site to have two alleles: a deleterious allele, M, and a neutral allele, A. We assume a Wright-Fisher co-dominance model to obtain:

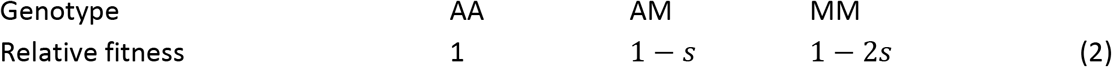

Following this model, we use the diffusion approximation 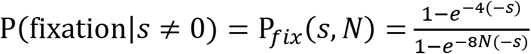 (48). Let P(*s*) describe the DFE of deleterious mutations, whose effects we assume to be independent of the environment. Substituting these into Eq.

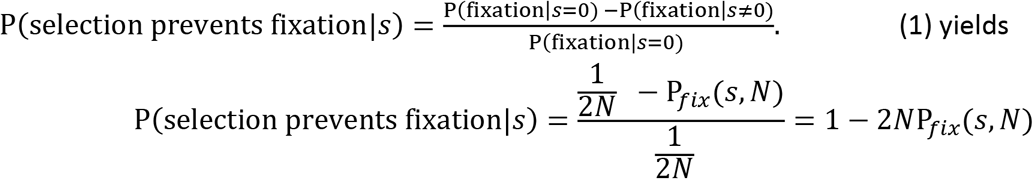

and

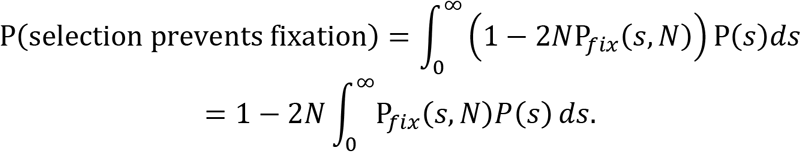

### Partitioning variance in log fitness by allele frequency

We model the deleterious allele M as having *j* copies, making its frequency 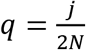 and the frequency of the allele A 1 − *q*. We assume random mating to obtain:

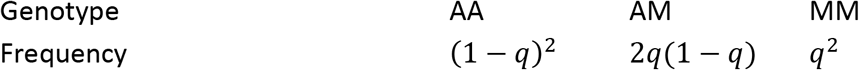

Let *r*_*i*_|*q* be a random variable specifying the contribution of a single site *i* with allele frequency *q* to the population’s natural logarithm of relative fitness *r. r*_*i*_|*q* can take the value log(1 − 2(*s*|*q*))^2^, log(1 − (*s*|*q*)), or 0, where *s*|*q* is also a random variable. Within-population variance in *r* contributed by this site is a random variable 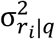 specified by (See the Supplementary section Per-site variance in log fitness):

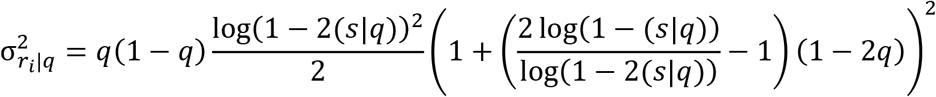

When *s* is small enough for log(1 − *s*) ∼ − *s*, this converges to a simpler expression for co-dominance on *r*, 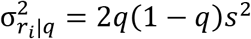 (derived in Supplementary Eq. S2), although here we use the full expression from above.

We next calculate the expected variance in log fitness 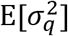 across sites with allele frequency *q* given that new mutations have a DFE, P(*s*), of *s*. Ignoring stochastic variation, a typical population will harbor *G* × P(*q*) sites with allele frequency *q*. Assuming independent evolution across sites, and using Bayes’ theorem P(*s*|*q*) = P(*q*|*s*)P(*s*) / P(*q*), yields

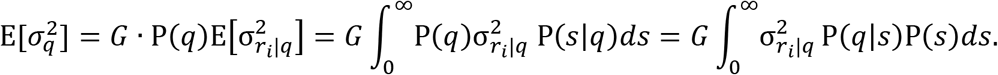

Many of the *G*_*del*_ sites outside of ancestral repeat regions are only partly constrained, meaning that some but not all mutations to them are deleterious. Our model simplifies this by considering the scenario with the same genome-wide deleterious mutation rate *U*, occurring at fewer sites *G*, at which all mutations are deleterious. This leads to a trivially small underestimation of variance because it allows two deleterious mutations to collectively rather than individually fix. We derive *G* from *U* = 2*μG*, using an estimated human per-site mutation rate *μ* = 1.515 × 10^−8^ as discussed in the Introduction. In other words, we assume that within the *G*_*del*_ sites outside of ancestral repeat regions, only *G* sites are constrained, and that a single conserved nucleotide identity is favored at each of these constrained sites.

Summing across all frequencies of the deleterious allele, the expected total variance in log fitness attributable to deleterious load is

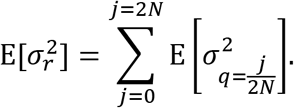

We estimate how much of the variance in load comes from deleterious alleles at frequency *q* as

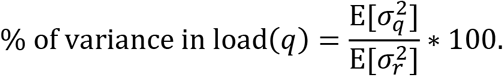

### *P*(*q*|*s*) under mutation-selection-drift

Under co-dominance, a diffusion model of bidirectional mutation, selection, and drift gives 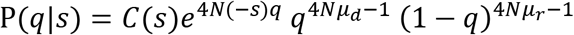 (49) for 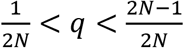 where *C*(*s*) is a normalizing constant, and *μ*_*d*_ and *μ*_*r*_ are the per-site deleterious and reversal mutation rates. A single preferred nucleotide at each of *G* sites corresponds to *μ*_*d*_ = *μ*, and 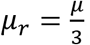.

Unfortunately, the diffusion approximation cannot capture the high probability, given low *μ*, that a site is monomorphic (i.e., P(*q* = 0)+P(*q* = 2*N*|*s*)) ((50) pp. 176-180).

To capture this probability using Queue theory (51), we assume that new mutations appear according to a Poisson process with rate 2*Nμ*. Each mutation spends time segregating in the population before getting lost or becoming fixed, with mean duration 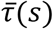. Then the per-site number of derived allele copies is Poisson distributed with mean 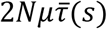, and the probability that a site is monomorphic is (Equation 6.22 in (51))

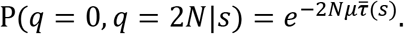

In our co-dominant Wright-Fisher model (Equation 1.62 in (50))

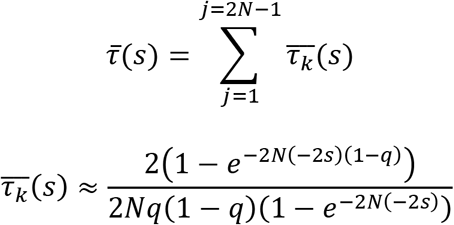

where 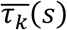 is the mean time that a new mutation spends at allele frequency 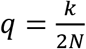 before fixation or loss.

## Results

*U* is larger than *U*_*obs*_ because it includes many weakly deleterious mutations that do not affect divergence data. *U* = 1.34U_*obs*_ (Figure 1A, orange) when we use an upper-bound heterozygote selection coefficient 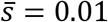 and CV=2.43 (see Introduction). Using the lower bound 0.00064 (See Introduction) produces *U* = 1.68U_*obs*_ (Figure 1A, green). The unweighted average of these two, 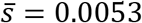 (Figure 1, blue), behaves much closer to the upper bound than to the lower bound. Thus, we combine *U* = 1.34U_*obs*_ with our bounds on human estimate 9.73 > *U*_*obs*_ > 2.85 (see Introduction) to yield 13 > *U* > 3.8.

**Figure 1.**
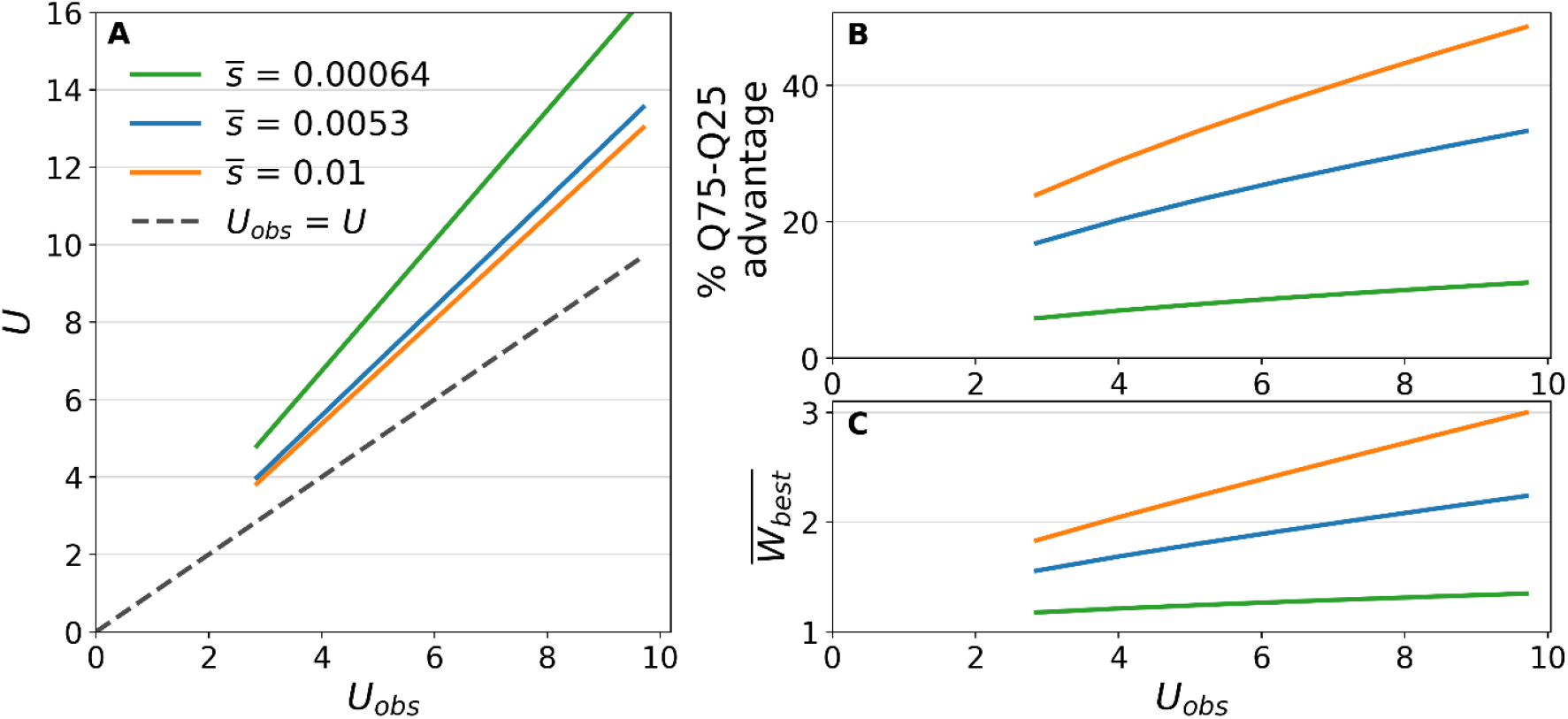
High genetic variation in fitness in ancient human populations under mutation-selection-drift balance does not exceed plausible reproductive limits. For all three DFEs, we used a coefficient of variation = 2.43 and *N* = 11,300 as inferred for nonsynonymous mutations (see Introduction). We used *μ* = 1.515 × 10^−8^ (see Introduction). Calculations are available in the Supplementary section Code figure 1.

To make variation in unconditional deleterious mutation load interpretable, we use the relative fitness advantage of a 75^th^ percentile individual over a 25^th^ percentile individual in the ancestral environment:

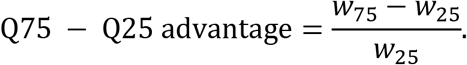

Note that this fitness advantage is quantified with respect to ancestral, not contemporary, human conditions, on the timescale that shaped the site frequency spectra from which the DFE was inferred. Fitness differences are nevertheless of interest because of the association between unconditional deleterious load and disease susceptibility (See Introduction).

The Q75-Q25 advantage is 24 − 49% (Figure 1B, orange) when using our upper bound 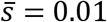 across the plausible range for *U*_*obs*_. This decreases to 6 − 11% (Figure 1B, green) when we use our lower bound 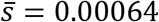. Using the average 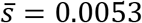 (Figure 1B, blue) gives a Q75-Q25 advantage of 17 − 33%.

Variance in load is not so high as to create a paradox with respect to the limits of human life history. 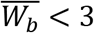 for all parameters tested (Figure 1C). This is lower than the human upper bound 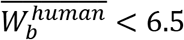 (33).

Given that the whole-genome DFE is a mixture between site classes, we expect the coefficient of variation to be larger than inferred for a single class. Increasing it from 2.43 to 4 yields substantially larger values of *U*, Q75-Q25 advantage and 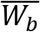, but a more reasonable increase to 3 has a more modest effect (Fig. S1).

In agreement with (47), rare mutations (<1% frequency, Figure 2A shading) explain most of the variation in human ancestral load. They explain 99% of the variance in log fitness when using our upper bound of 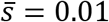 (Figure 2A, orange at the right edge of shading), and 83% when using our lower bound of 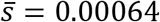 (Figure 2A, green at the right edge of shading). Most of the variance in log fitness is correspondingly explained by mutations with selection coefficients larger than the mean (Figure 2B, compare each cdf to the vertical line of matching color) and far larger than the drift barrier of 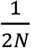 (Figure 2B, black vertical line). Again, a halfway value of 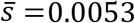, representing an average between the bounds for 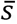, more closely resembles the upper bound.

**Figure 2.**
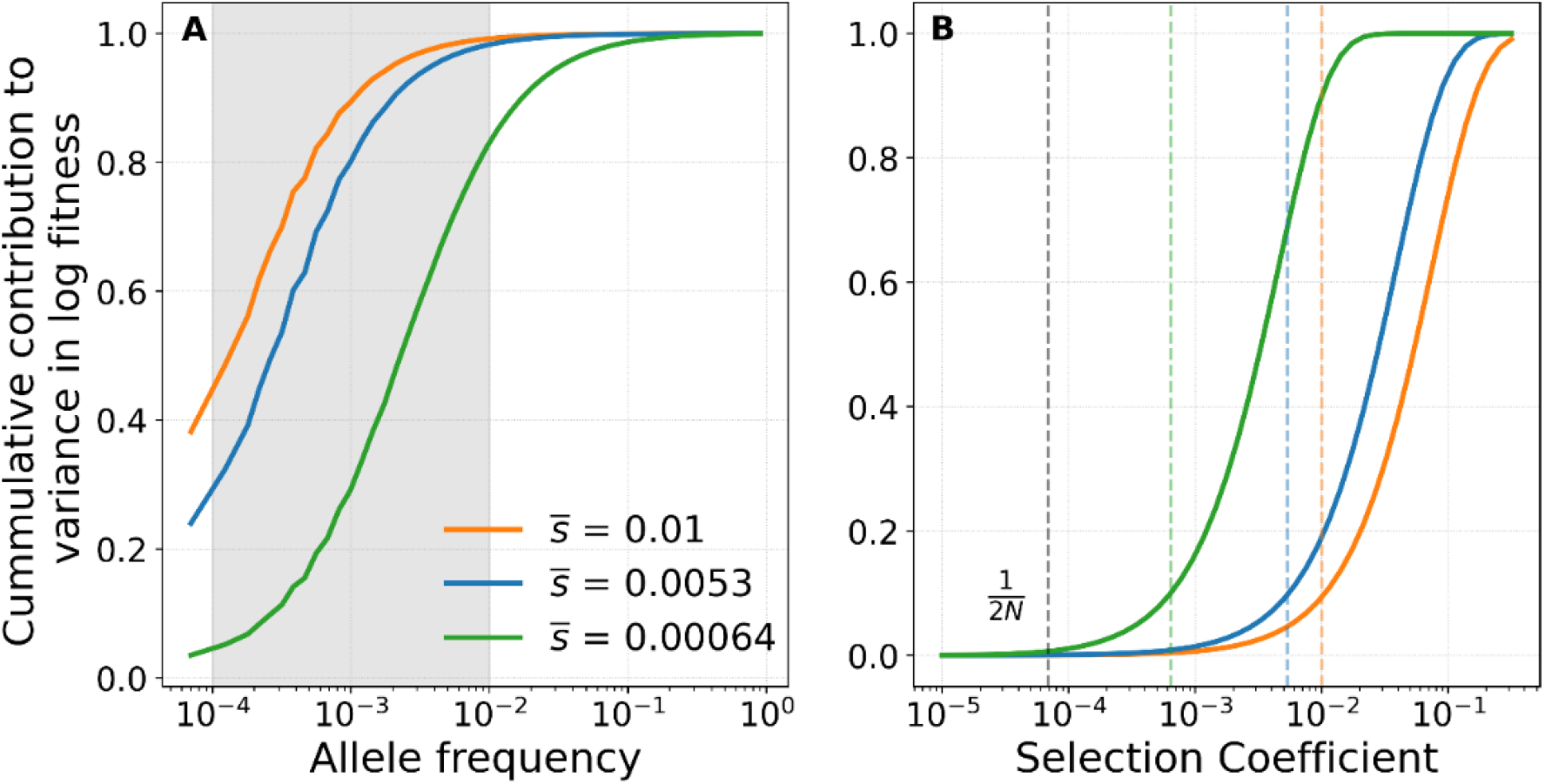
Most variation in ancestral human load under mutation-selection-drift balance comes from rare, strongly selected alleles. For all three DFEs, we used a coefficient of variation = 2.43 and *N* = 11,300 as inferred for nonsynonymous mutations (see Introduction). We used *μ* = 1.515 × 10^−8^ (see Introduction). 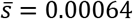 and 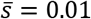 are the lower and upper human bounds, respectively (See Introduction). 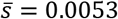 is a simple average of the two means. A) Gray area represents the rare alleles considered in (52) B) Vertical lines represent the drift barrier (gray), and the averages for each DFE (colored). Calculations available in the Supplementary sections Code figure 2 (1) and Code figure 2 (2).

Drift reduces the variation in load and 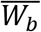 relative to values expected from deterministic mutation-selection balance (Figure 3). (24) previously suggested the opposite in models that neglected the probability that sites are monomorphic and focused on lag load rather than load variance.

**Figure 3.**
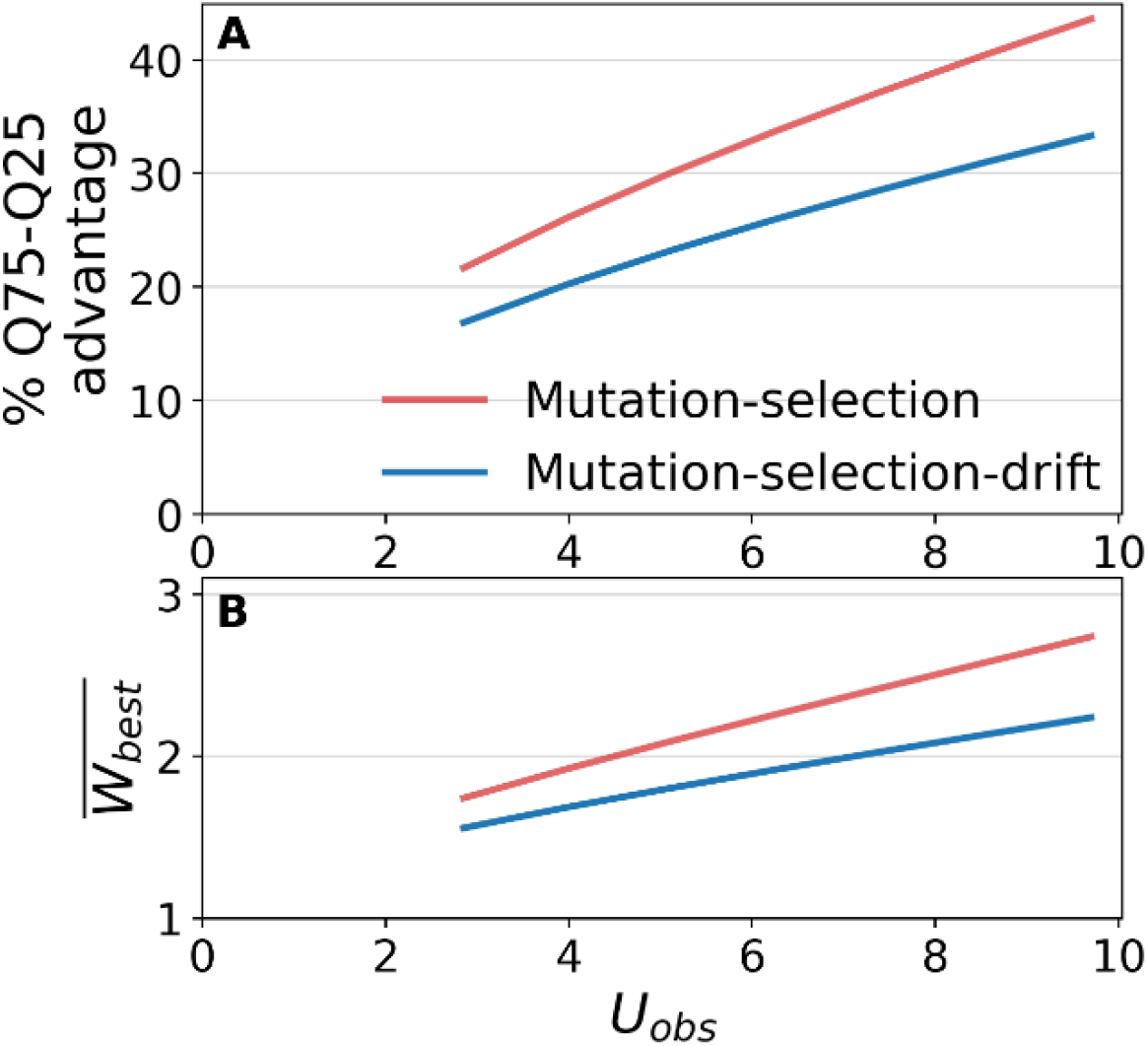
Drift modestly decreases variation in load. Variance in log relative 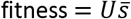 for mutation-selection balance (See Introduction). For mutation-selection-drift balance, we used a DFE with coefficient of variation = 2.43 and *N* = 11,300 as inferred for nonsynonymous mutations (see Introduction). In both models, we used our intermediate mean selection coefficient 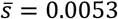, and *μ* = 1.515 × 10^−8^ (see Introduction). Calculations available in the Supplementary section Code figure 3.

High *U* is widespread across vertebrates (37, 38, 53), suggesting that high load variation is not restricted to humans. In Figure 4, we explore a range of *N* values (in our Wright-Fisher model) consistent with coalescent times as low as the vaquita, *N*_*e*_ = 3500 (54), up to as high as the brown rat, *N*_*e*_ = 124000 (55). Non-synonymous mutations’ 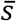 might vary by two orders of magnitude across vertebrates (56). Given even greater uncertainty in the whole genome’s DFE, we vary 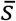 by three orders of magnitude on the x-axis of Figure 4, while keeping the coefficient of variation constant.

**Figure 4.**
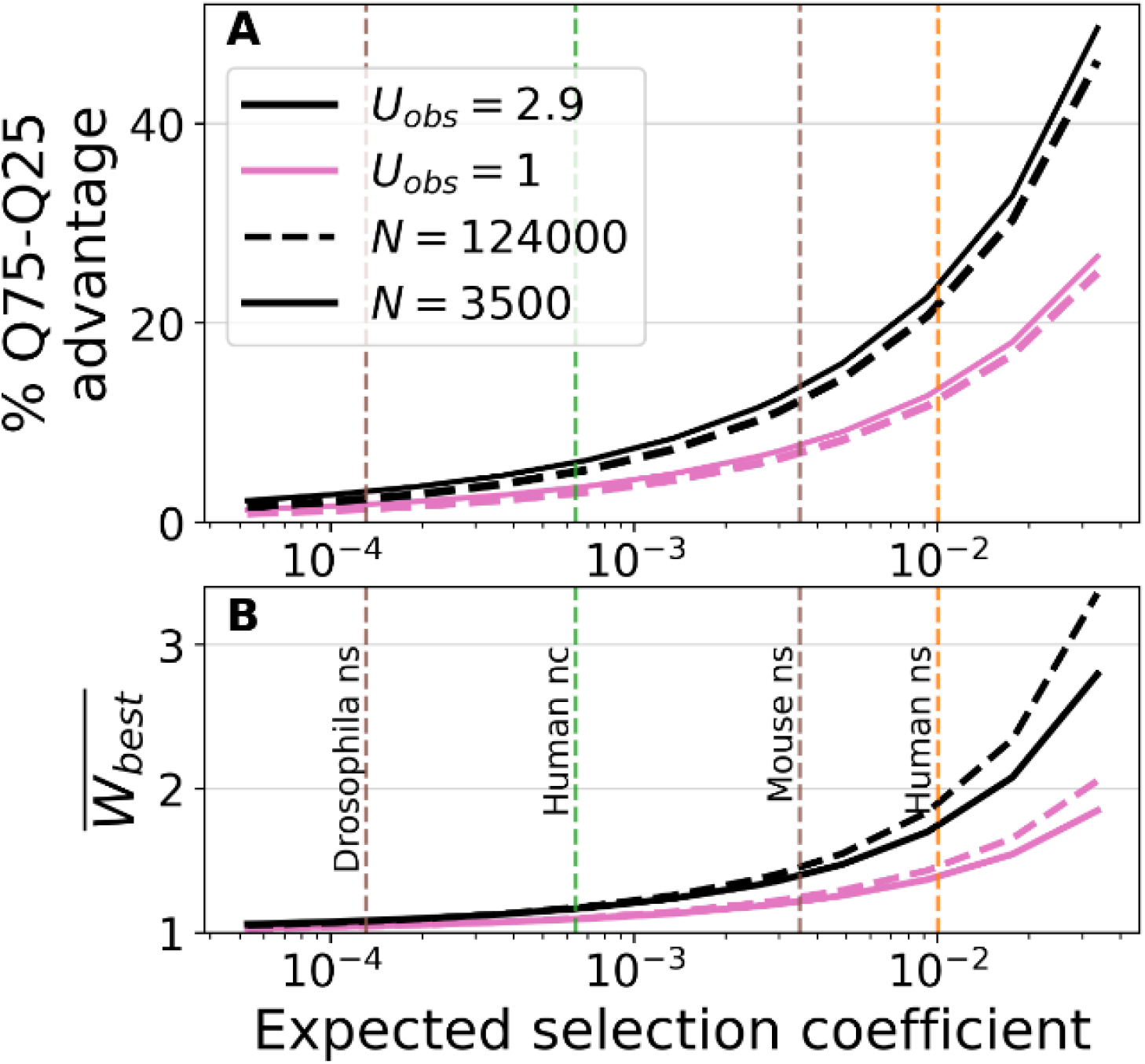
High variation in load depends on strong selection, but not on low *N*_*e*_. The pink lines represent the Mouse’s lower bound *U*_*obs*_ > 1 (37). We calculate an upper bound (*U*_*obs*_ < 2.9, black lines) by combining *G*_*cons*_ = 9% of the 2.5 × 10^9^ mouse diploid genome sites (44), and a per-base mutation rate, *μ* = 6.53 × 10^−9^. The latter combines indels and point mutations (42). For Drosophila, we calculate an upper bound *U*_*obs*_ by assuming that all mutations arriving on any of its 1.4 × 10^8^ diploid sites, at a rate *μ* = 5.45 × 10^−9^ (42) are deleterious.This yields *U*_*obs*_ < 1.5. Drosophila and mouse estimates of 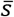 for nonsynonymous mutations (ns) were inferred in (56). Human nc corresponds to 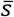 for mutations at ultra-conserved noncoding sites (See Introduction). We use Mouse’s coefficient of variation = 2.13 and *μ* = 6.53 × 10^−9^ throughout the plot. Calculations available in the Supplementary section Code figure 4.

High variation in load depends on large 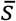 and *U*. A mammal such as Mouse meets these criteria. (56) inferred the mouse non-synonymous DFE as a gamma distribution with mean 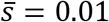 and coefficient of variance CV=2.13. Combining this DFE with 2.9 > *U*_*obs*_ > 1 (see Figure 4 caption) results in a 6-12% Q75-Q25 advantage (Figure 4A). In contrast, Drosophila’s non-synonymous 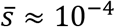 and CV = 1.69 (56) makes the Q75-Q25 advantage smaller than 3% (Figure 4A) even under an unrealistically high *U*_*obs*_ = 2.9 (see Figure 4 caption) and inflated CV=2.13 (which inflates variation in load (Fig. S1B)). In contrast to the importance of 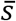 and *U*, varying population size, which affects the drift component of mutation-selection-drift-balance, does not qualitatively affect the variation in load (Figure 4, dashed vs solid lines). 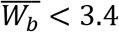 for the entire range of parameters examined in Fig 4B, including unrealistically large whole-genome 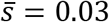. This does not pose a paradox, since most species can easily accommodate reproductive excess higher than 4.

## Discussion

Our results suggest that each human newborn carries, on average, more than 4 new deleterious mutations. This high mutational input leads to high variation in human deleterious load, corresponding to a 17-33% difference in fitness between two typical individuals in the human ancestral environment. This high load variation is consistent with population persistence, since our model’s best-fitness individual 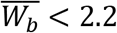 is consistent with human life history traits. High load variation is driven primarily by rare variants. Our results suggest that high variation in deleterious mutation load will be found in any species with high 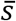 and high *U*.

Our 17-33% range for a typical fitness difference reflects uncertainty in *U*. Also considering uncertainty in 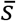 broadens the range to 6-49%. We note that (26) argued that variance in load is low, but assumed unrealistically small selection coefficients near the drift barrier of *s* = 1/2*N*. Positive global epistasis is expected to increase fitness variance above that in our non-epistatic model, while negative global epistasis will decrease it (22, 57, 58). However, realistically weak global epistasis (59–61) has a negligible effect on the Q75-Q25 spread in load (62).

Here, we solved for mutation-selection-drift balance, in contrast to (36)’s solution for mutation-selection balance. (24) claimed that drift increases mutation load, using a model in which allele frequency could drift, but alleles were never lost or fixed, an assumption that requires unrealistically high per-site mutation rates. In contrast, our mutation-selection-drift balance solution incorporates the probability that a site is monomorphic. Our results imply lower variance in load and 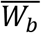 than either of these two previous models (Figure 3). However, we still judge variance to be substantial when we consider more realistic values for human *U* and 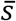 than they did.

Some genetic variation in fitness might arise from sources other than unconditionally deleterious mutations, particularly balancing selection (63–65). Empirical studies in *Drosophila* suggest that, for most traits, unconditionally deleterious variation accounts for only a small proportion of the genetic variation (63, 65). However, the theory presented here expects mammals to be different to *Drosophila*, with a substantial role for unconditionally deleterious load requiring large mean selection coefficients and large *U*, which mammals but not *Drosophila* have.

Variance in fitness is important for a variety of phenomena, the most obvious being the rate of genetic adaptation (45, 66). Selection against low-fitness individuals creates background selection and linkage disequilibria even among unlinked sites (67, 68). Differential effects of background selection on rare vs. common alleles can distort the site frequency spectrum (69, 70), and have complex effects on trait variation (70). Limits to 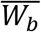 create a maximum size for the functional genome (36), and for the number of traits under selection (71).

Our finding of high variation in human load is consistent with the omnigenic model (72). This model was proposed to explain why disease-associated genetic variants are distributed broadly across the genome rather than concentrated in key molecular pathways. In this model, some ‘core genes’ are directly associated with disease, while many other ‘peripheral genes’ are indirectly associated through their regulation of core genes. Because of the importance of peripheral genes, much disease variation is distributed among many variants of small effect (71, 73, 74). Genetic load can be thought of as mutations to peripheral genes that affect many different diseases. A deleterious mutation of moderately large selective effect might still have only small effects on each individual disease. In agreement with this view, rare, strongly deleterious mutations have larger effects on overall gene expression (75), and affect more cell types than common mutations (76). More directly, an individual’s load in coding regions, as assessed by the number and conservation scores of mutations, is associated with the presence of at least one disease (77).

Our results suggest that rare variants are an important component of disease susceptibility. GWAS studies that incorporate whole-genome sequencing can capture rarer variants, establishing their importance to disease (71, 74). (52) estimated that, on average across 34 complex traits and diseases, alleles with frequencies between 0.01% and 1% (Figure 2A, grey shading) explain around 20% of the genetic variance estimated from pedigrees. In the more extreme case of height, including whole-genome sequencing increases the heritability explained by single-nucleotide polymorphisms from 48% to 68%, compared to a heritability of 70-80% estimated from pedigrees (78). Our Fig. 2a suggests that while the 0.01%- 1% range is the most important, variants at frequencies below 0.01% could also be substantial, especially if the mean selection coefficient is large.

Polygenic Risk Scores (PRSs) are a popular approach to identifying high-risk individuals (79). PRSs use summary statistics from GWAS to predict disease liability. The individuals most likely to benefit from early medical intervention are found at the tail of the liability distribution. Incorporating rare variants into the GWAS underlying a PRS improves identification of individuals at the extreme tail of disease risk more than it improves overall predictive power throughout the distribution (6). Variant effect prediction that bypasses GWAS can incorporate even rarer variants, resulting in superior identification of individuals at extremely high, clinically actionable risk (7).

Despite these recent advances, assessing the effect of rare variants on disease susceptibility remains challenging (80). For example, the statistical power of GWAS is lower for rare alleles (3, 5). Although annotating variants on the basis of additional information can increase the power of these tests (81, 82), annotating noncoding regions remains challenging (but see (76, 81, 83) for recent advances).

High human variation in genetic susceptibility to disease has been used to rationalize discriminatory policies (reviewed in (84)), including forced sterilization to reduce the number of children of ‘loaded’ individuals. These ideas culminated in horrific human rights violations, mostly targeting disabled communities (85), women, and indigenous and ethnic minority groups (86–88). We condemn this use, aligning instead with the view that a better understanding of genetic diversity, including load, can promote more equitable societies (89– 91).

Our results suggest that rare, unconditionally deleterious mutations make a substantial contribution to overall genetic load. Because load might impact many diseases, rare deleterious variants could also contribute substantially to variation in disease. If this is true, then new methods for identifying these variants should help identify individuals at high risk of disease. Future work on rare variant detection will be useful in testing the hypothesis that load is important, as well as having a direct impact on precision medicine.

## Acknowledgments

We thank Micaila Marcelle, Ryan Gutenkunst, David Enard, and Bruce Walsh for helpful discussions, and Cock Van Oosterhout for comments on the manuscript. We thank the John Templeton Foundation [62028] and the National Science Foundation (DEB-2333243) for funding.

## Supplementary

## Supplementary Methods

### Per-site variance in log fitness

Let genotypes MM, AM, and AA have a trait value of 2*a, a*(1 + *k*), and 0, respectively. Under a diploid single-site, random mating model, the within-population genetic variance is (Equation 4.12a in (1)):

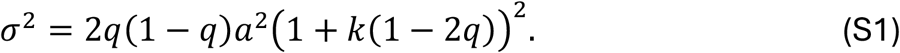

Using our per-generation fitnesses from Eq. (2) and considering the natural logarithm of relative fitness *r* as our focal trait, yields 2*a* = log(1 − 2*s*) and *a*(1 + *k*) = log(1 − *s*). Rearranging leads to 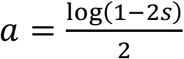 and 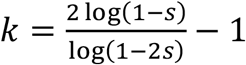. Substituting these two terms into Equation S1 yields

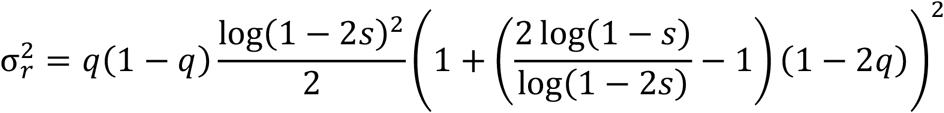

For typically small *s*, log(1 − *s*)∼ − *s*. This makes *r* a co-dominant trait where genotypes MM, AM, and AA have an *r* value of −2*s*, −*s*, and 0, respectively. Under this scenario

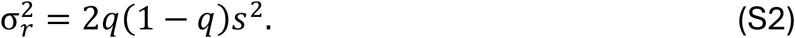

### Expressing fitness variation via quantiles

Assuming no epistasis nor linkage equilibrium, an individual’s log relative fitness *r* is equal to the sum of the contributions to log fitness across *G* sites. Since *G* is of order 10^8^ (see the Introduction), we use the Central Limit Theorem to approximate the distribution of *r* as Normally distributed. This makes relative fitness (*e*^*r*^) approximately log-Normally distributed. To make variation in load easily interpretable, we express results in terms of the Q75 − Q25 fitness advantage:

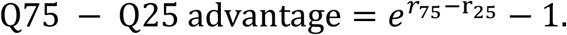

where *r*_*x*_ is the *x*^th^ percentile of log relative fitness. The quantile function of *r* is 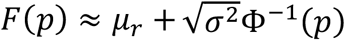, where Φ^−1^(*p*) is the *p*^th^-quantile of the standard normal distribution and *σ*^2^ is its variance. Substituting this into the equation above yields

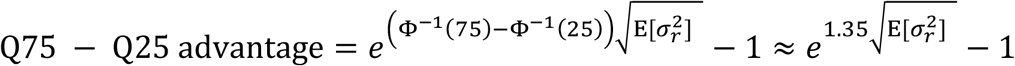

where we replace 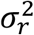 by its average E[*σ*^2^].

### Best individual’s absolute fitness

As discussed in the Introduction, to avoid a paradox, the best individual’s absolute fitness 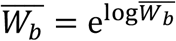 must be plausible for the species’ life history. Absolute fitness *W* is related to relative fitness (*e*^*r*^) via a normalizing constant *e*^*a*^. This yields

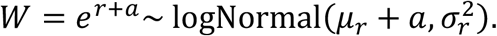

At demographic equilibrium, the average number of offspring per individual is 1, which is also the mean of a log-normal distribution, such that 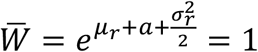. This means 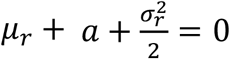. Substituting 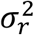 with its expectation 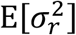 yields

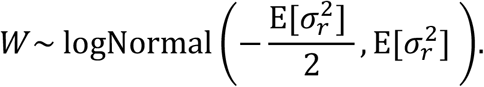

To estimate 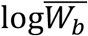, we assume linkage equilibrium and model a population as a random sample of size *N* from the distribution of 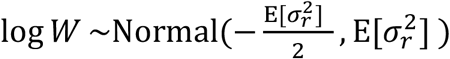. 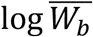 is then the expected maximum value of our sample. Using Extreme Value Theory (2)

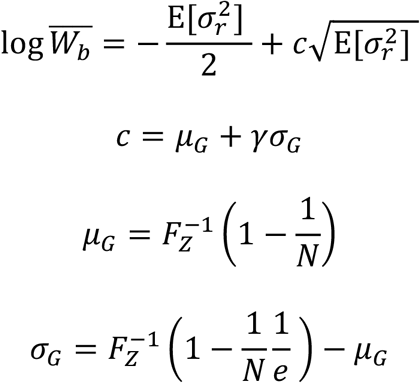

where *γ* is the Euler-Mascheroni constant, 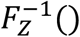 is the quantile function of a standard Normal distribution and *e* is the Euler number.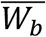 is then calculated as 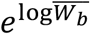.

## Supplementary Figure

**Figure S1.**
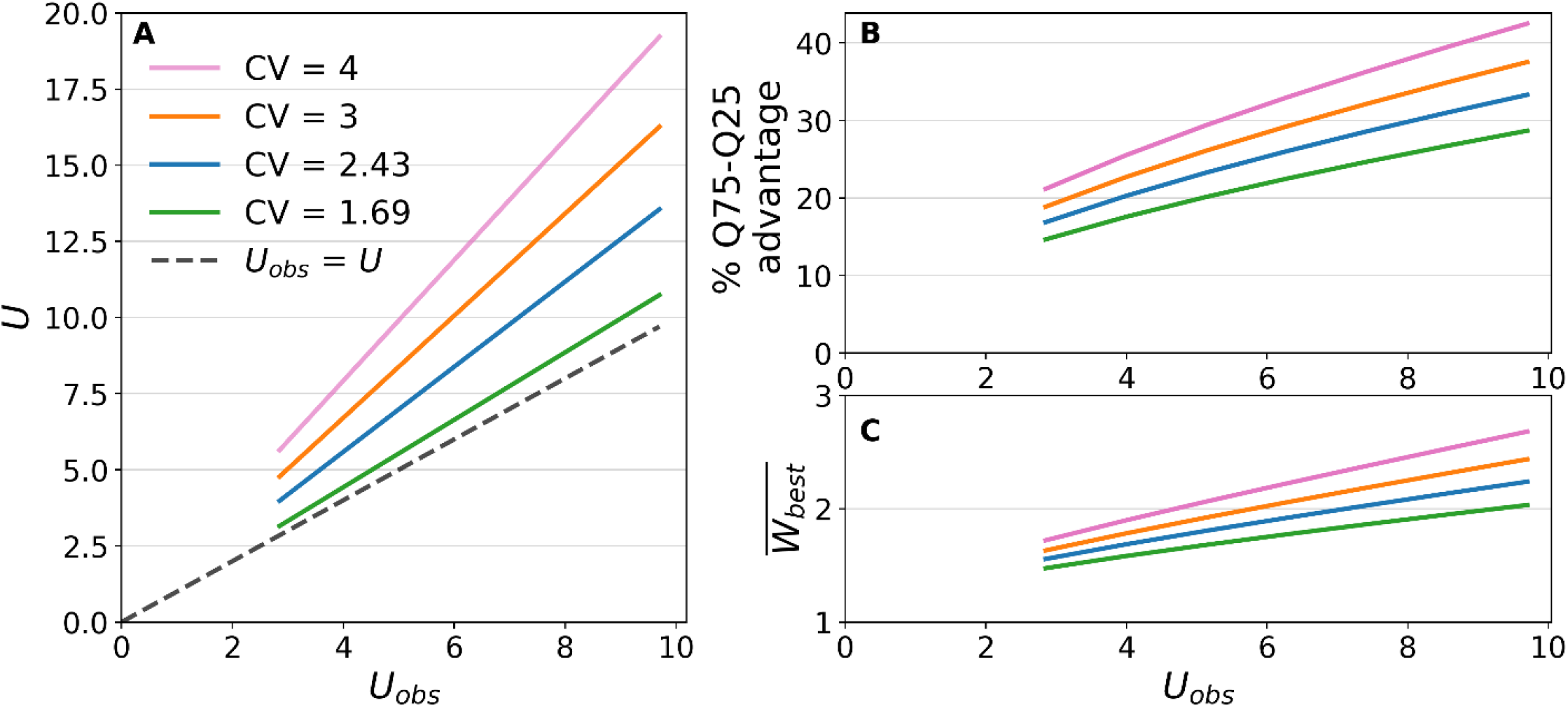
DFEs with a higher coefficient of variation (CV) produce higher variation in fitness. The green line is the inferred CV for Drosophila’s nonsynonymous mutations (3). Blue line is the default CV used in the main text inferred for nonsynonymous mutations (4). For all four DFEs, we used 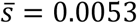 as an average between the upper bound and lower bound human 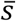 (see Main text). We used *N* = 11,300 as inferred for human nonsynonymous mutations (4), and *μ* = 1.515 × 10^−8^ (see Introduction).

---

**Code figure 1**

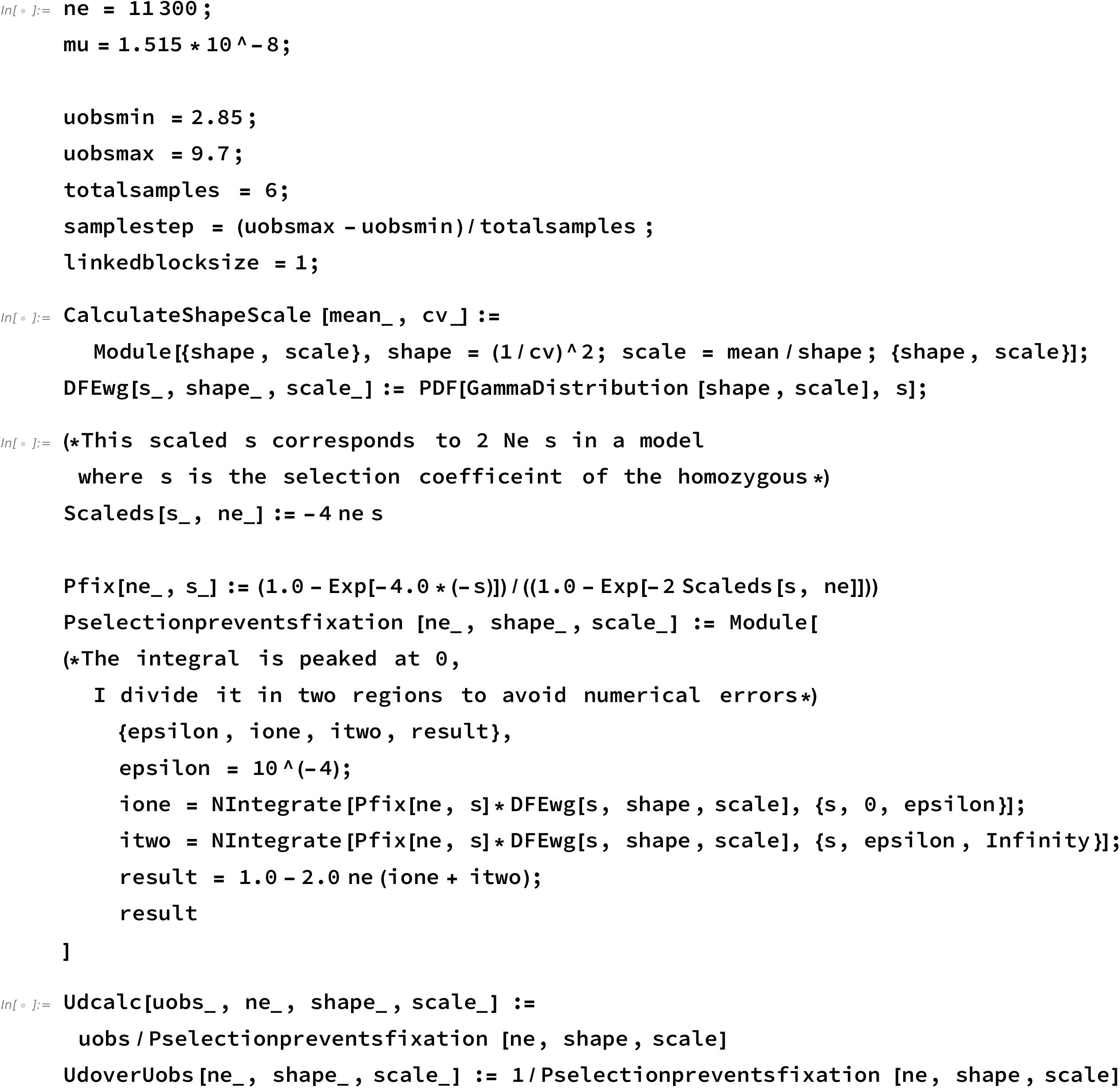
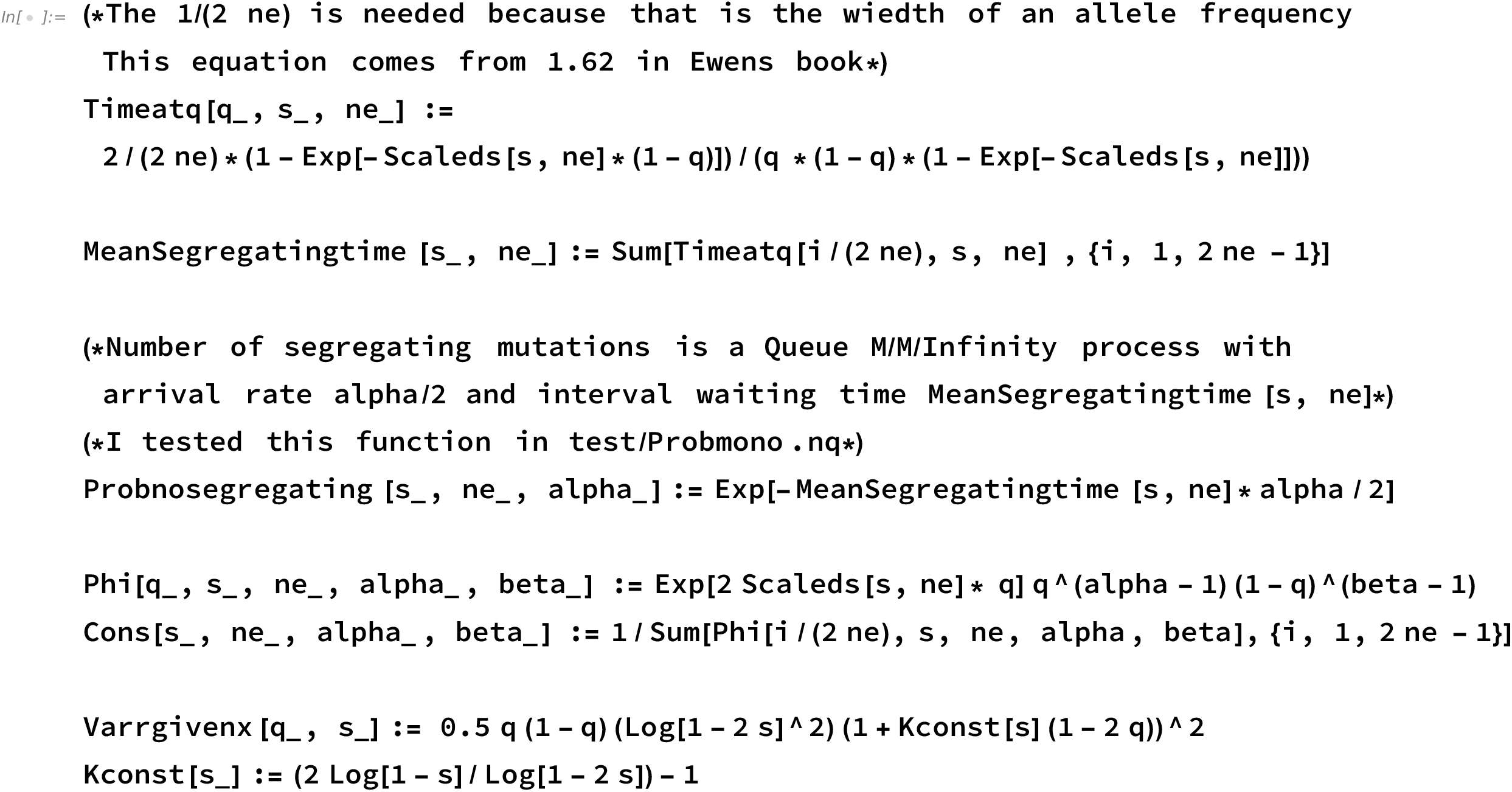
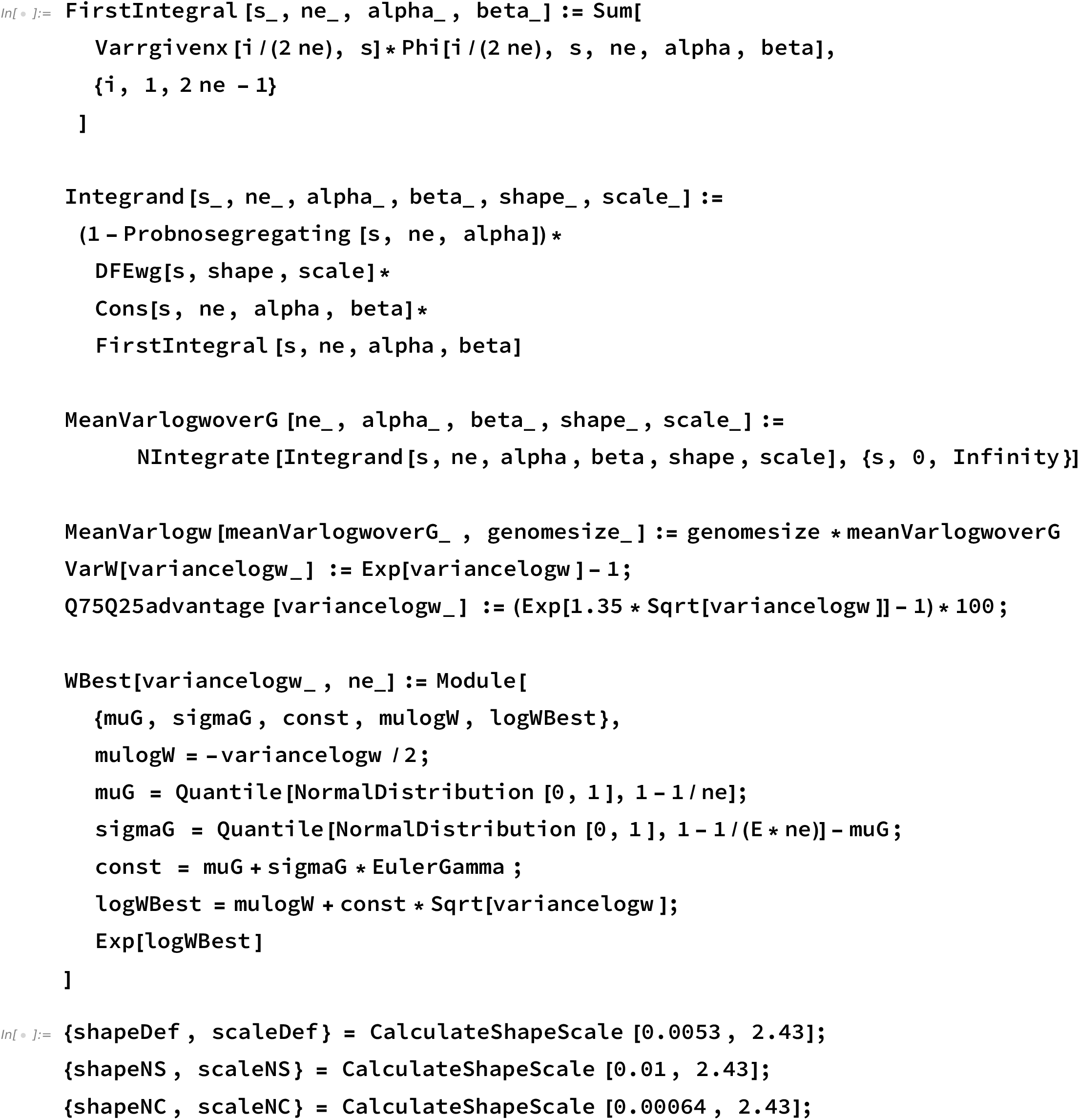
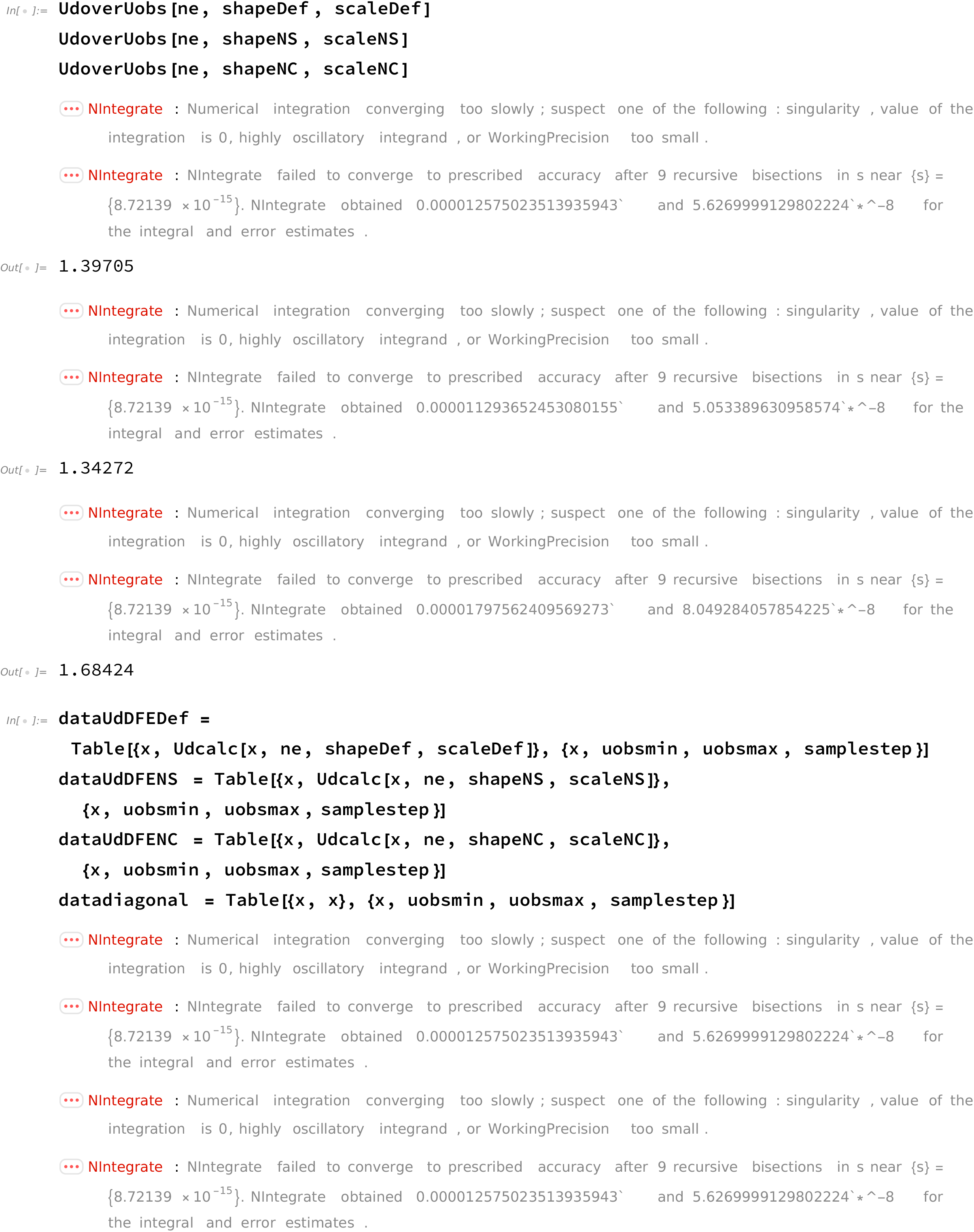
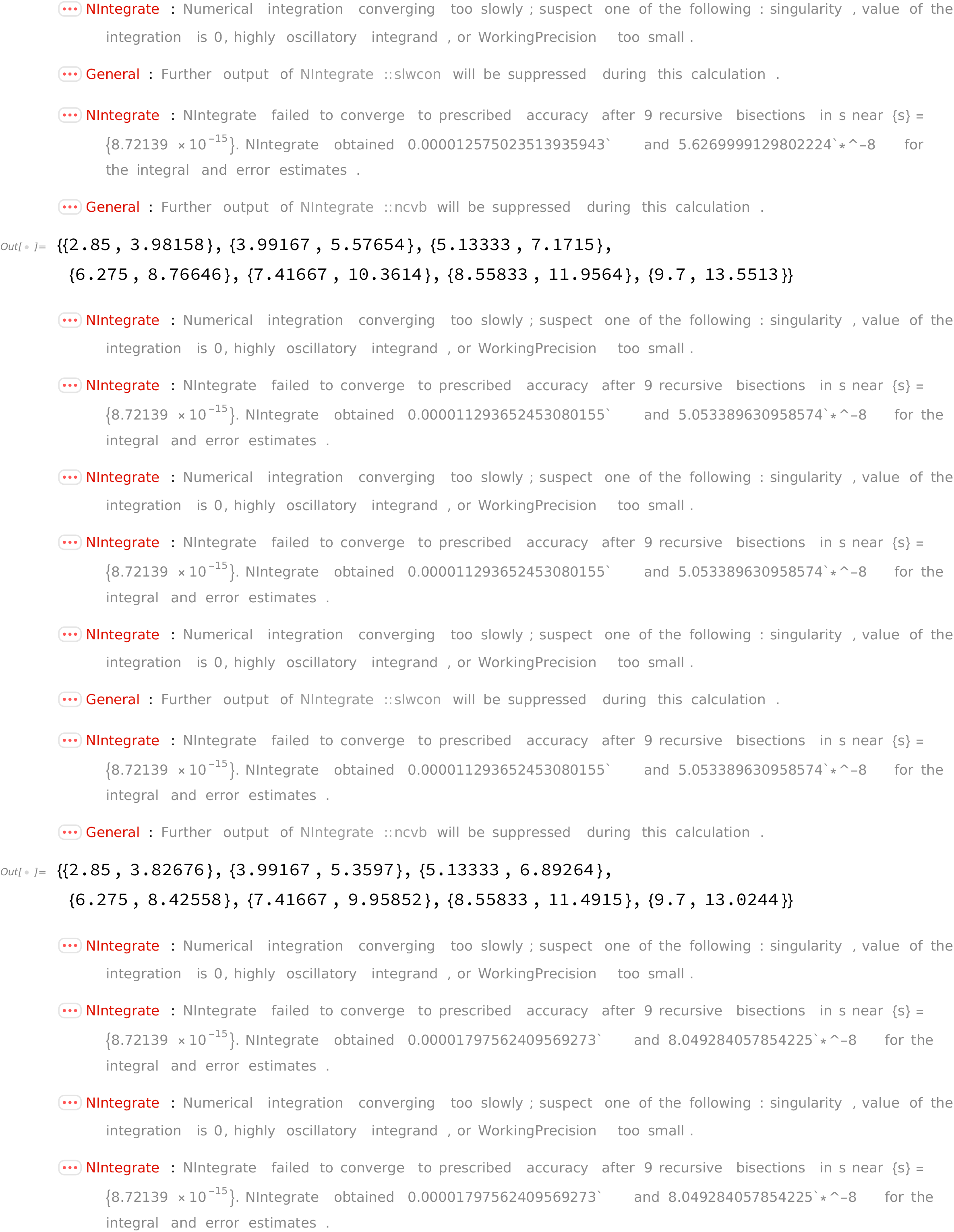
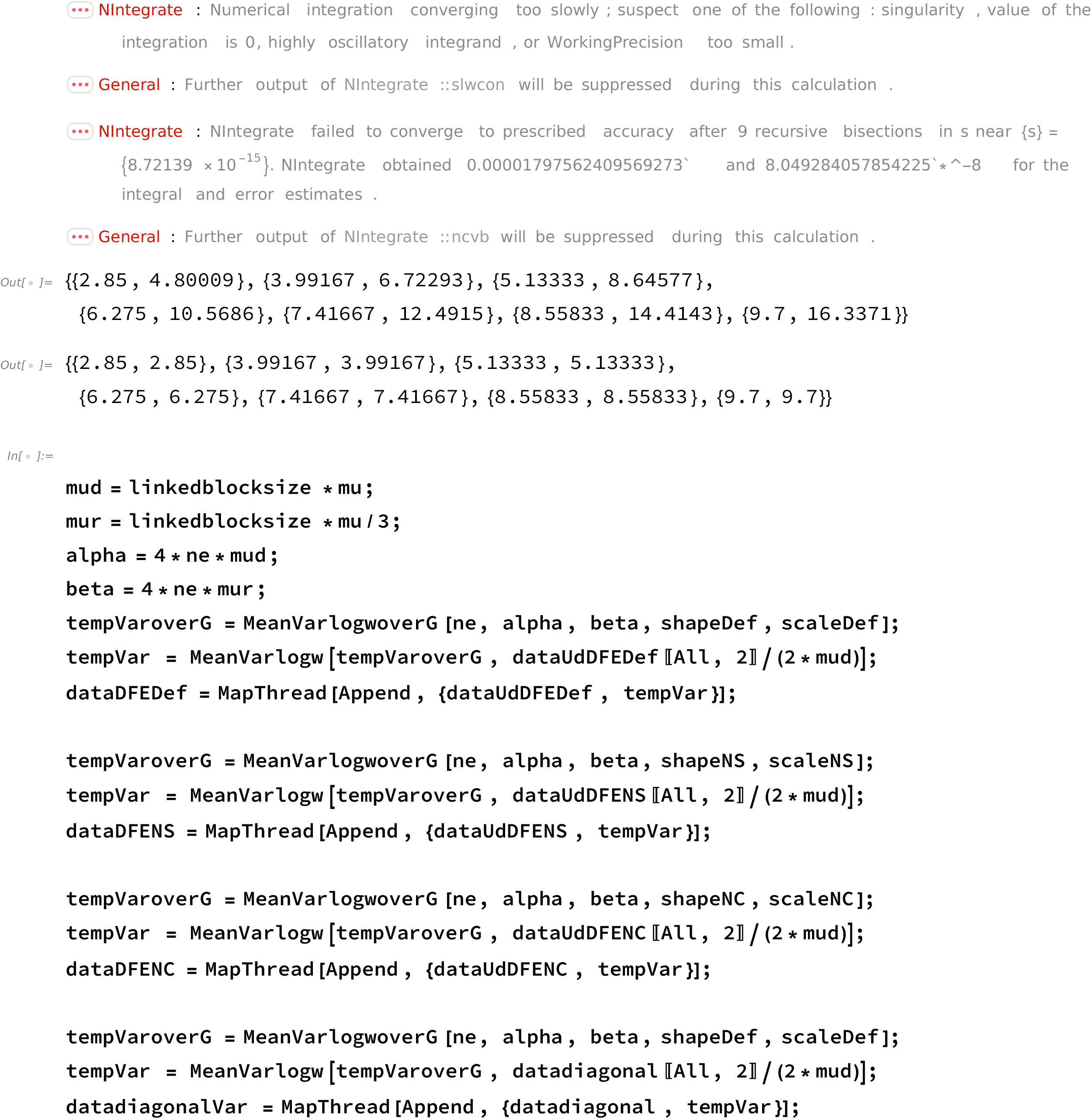
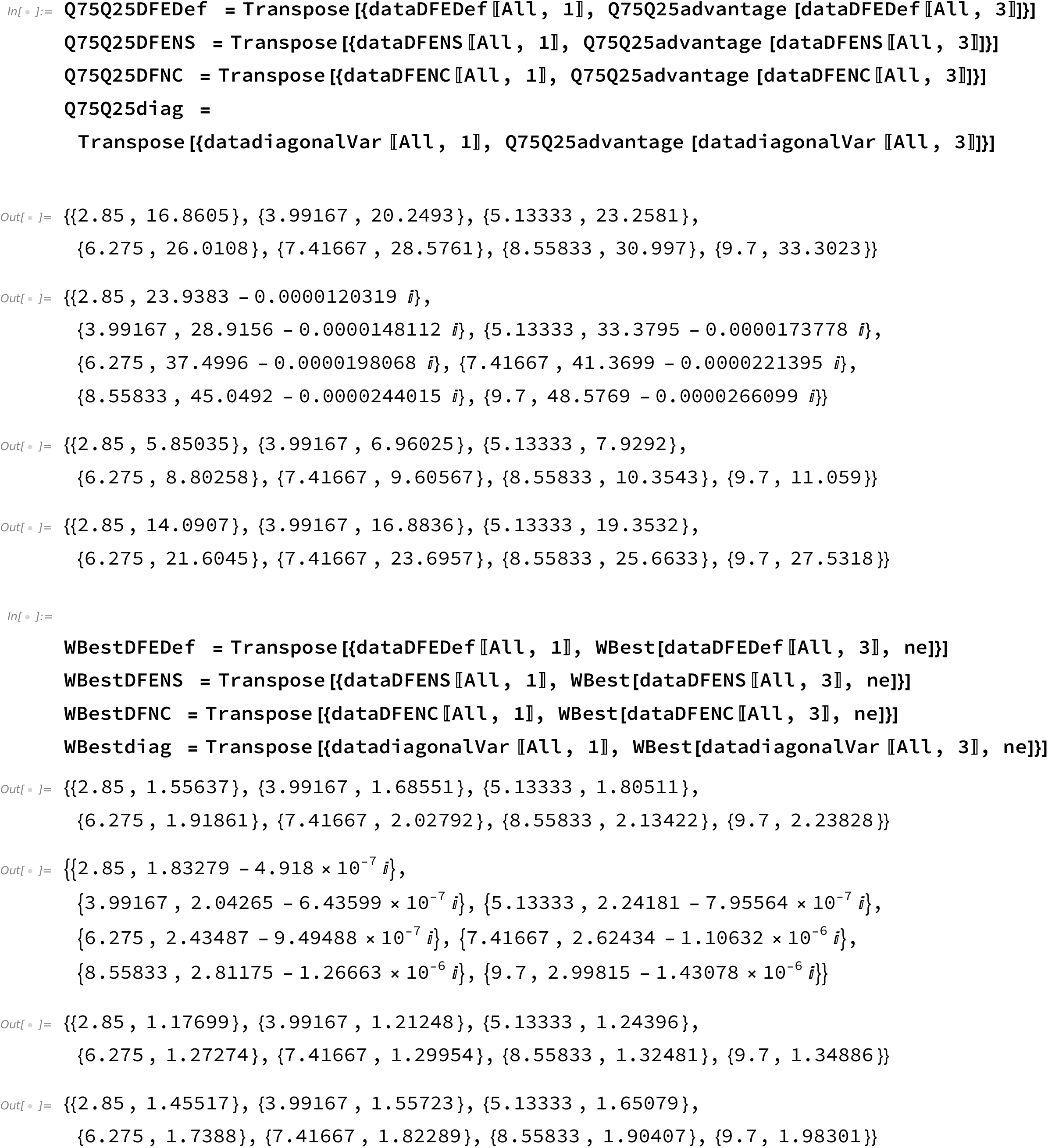

**Code figure 2 (1)**

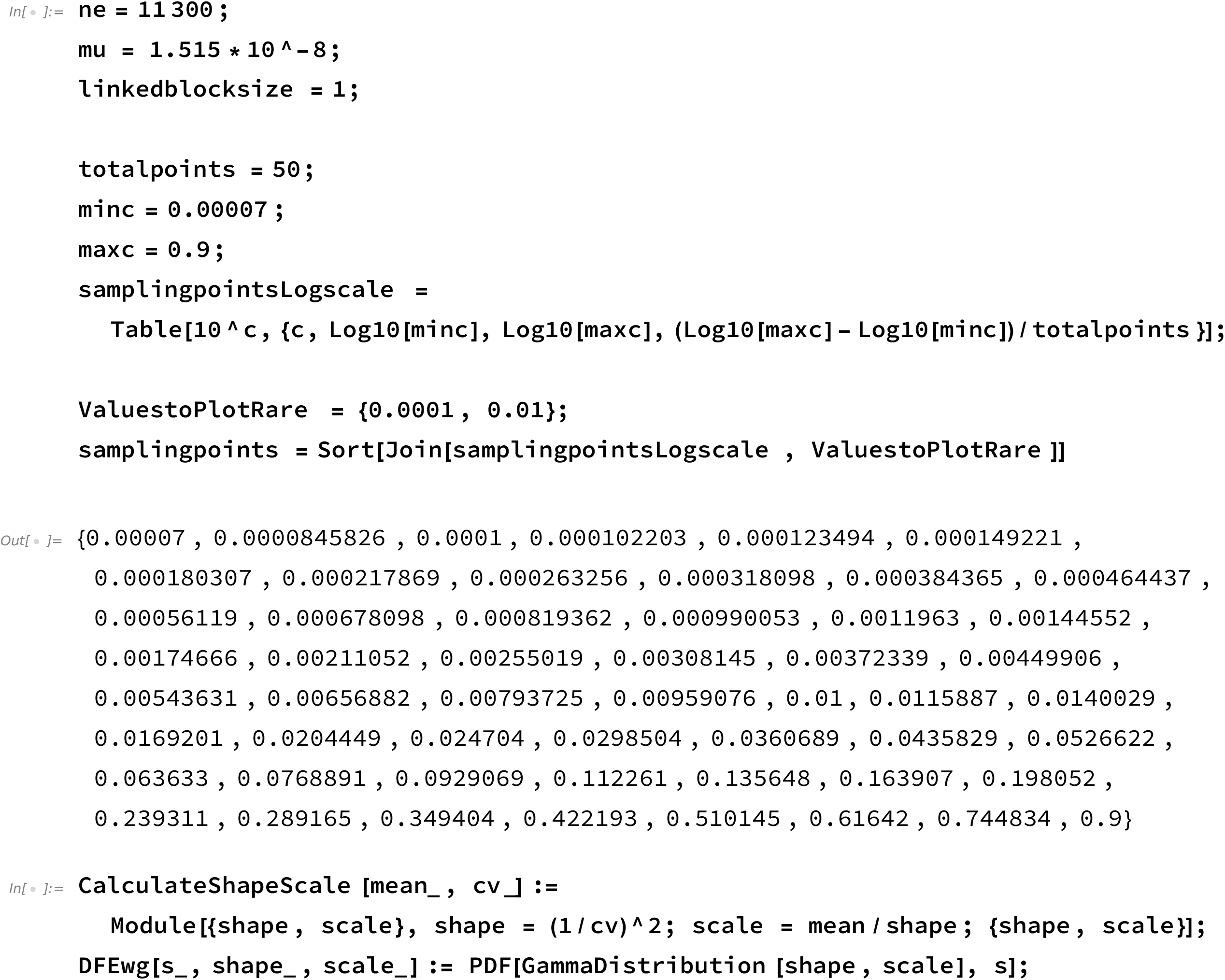
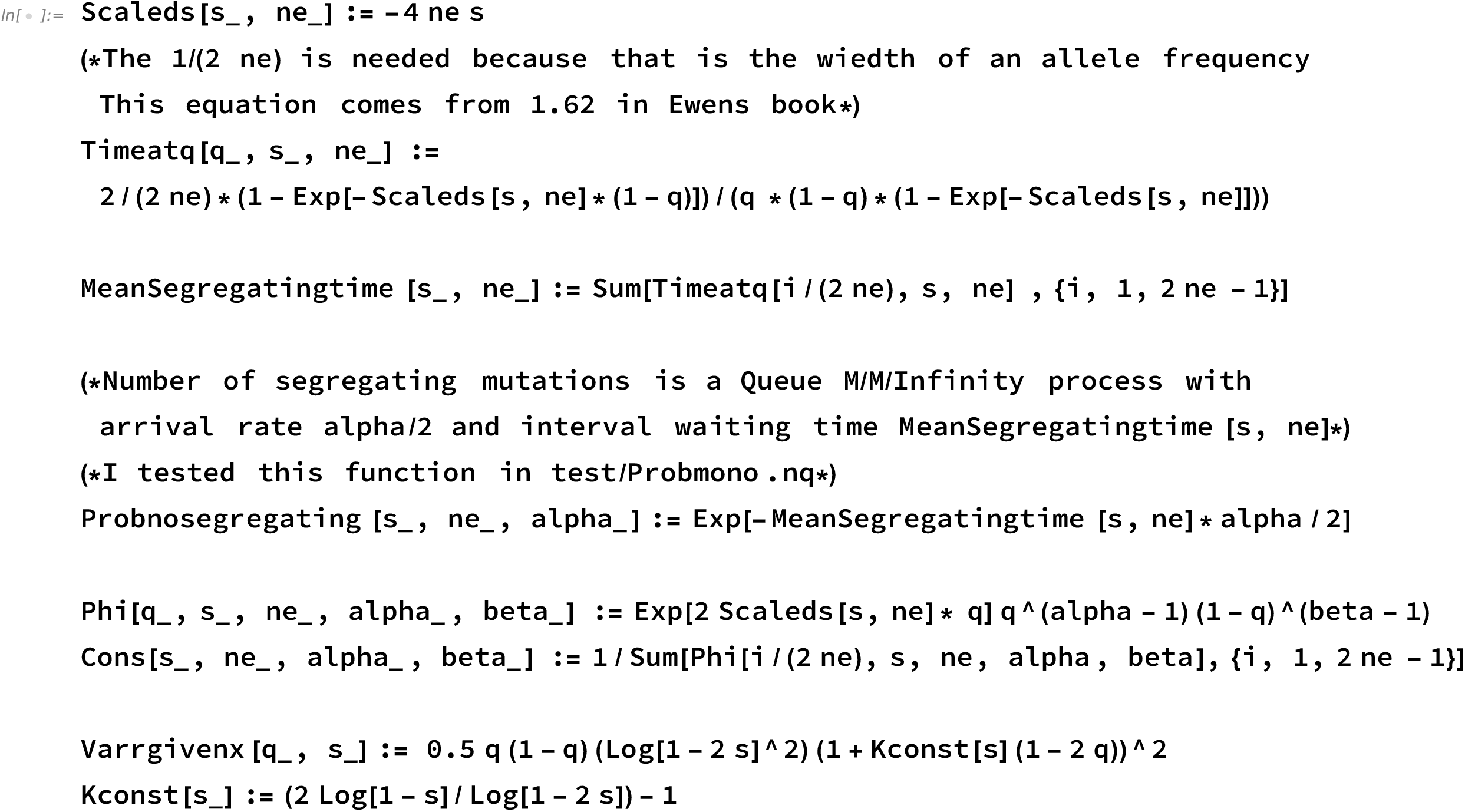
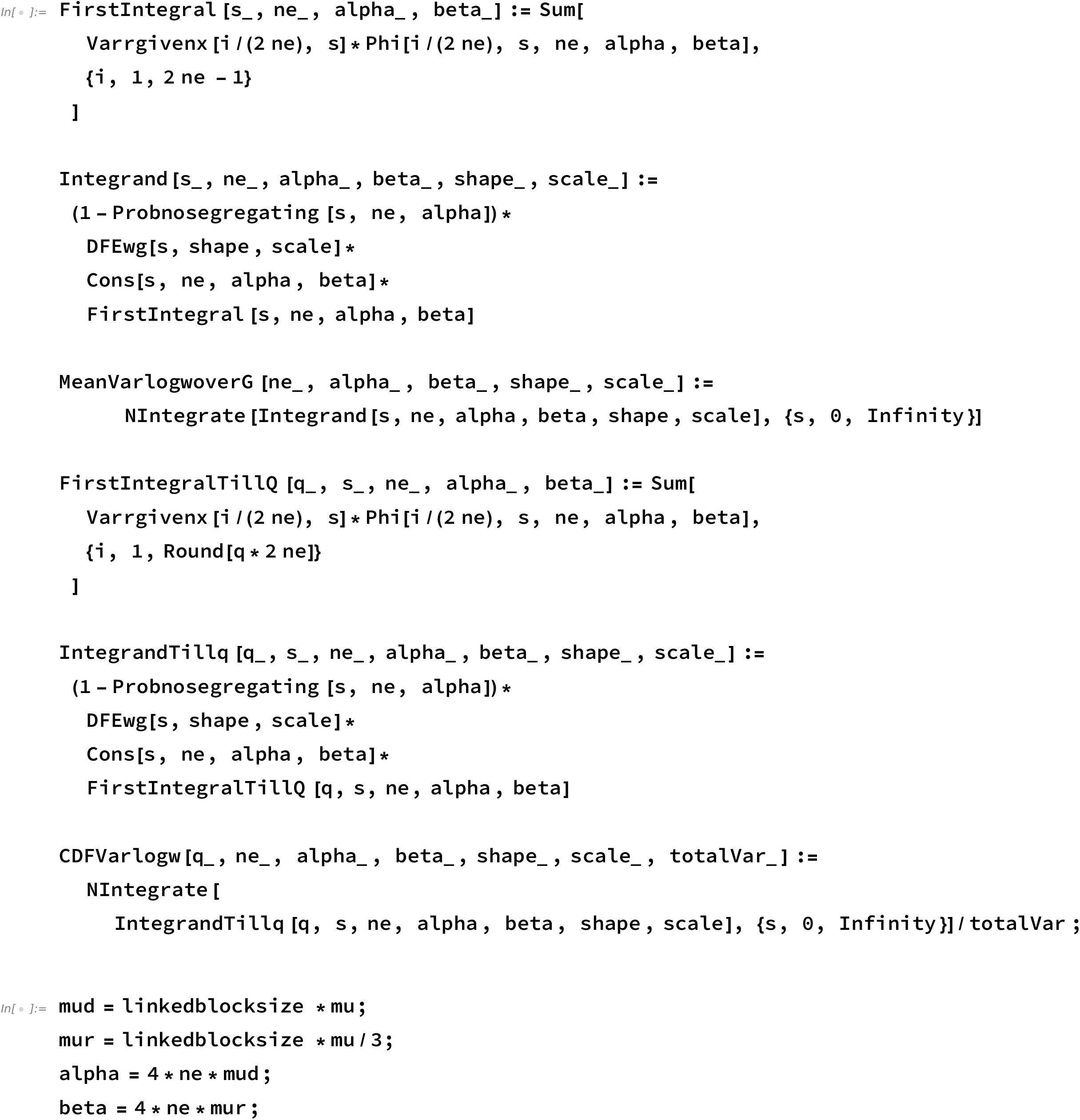
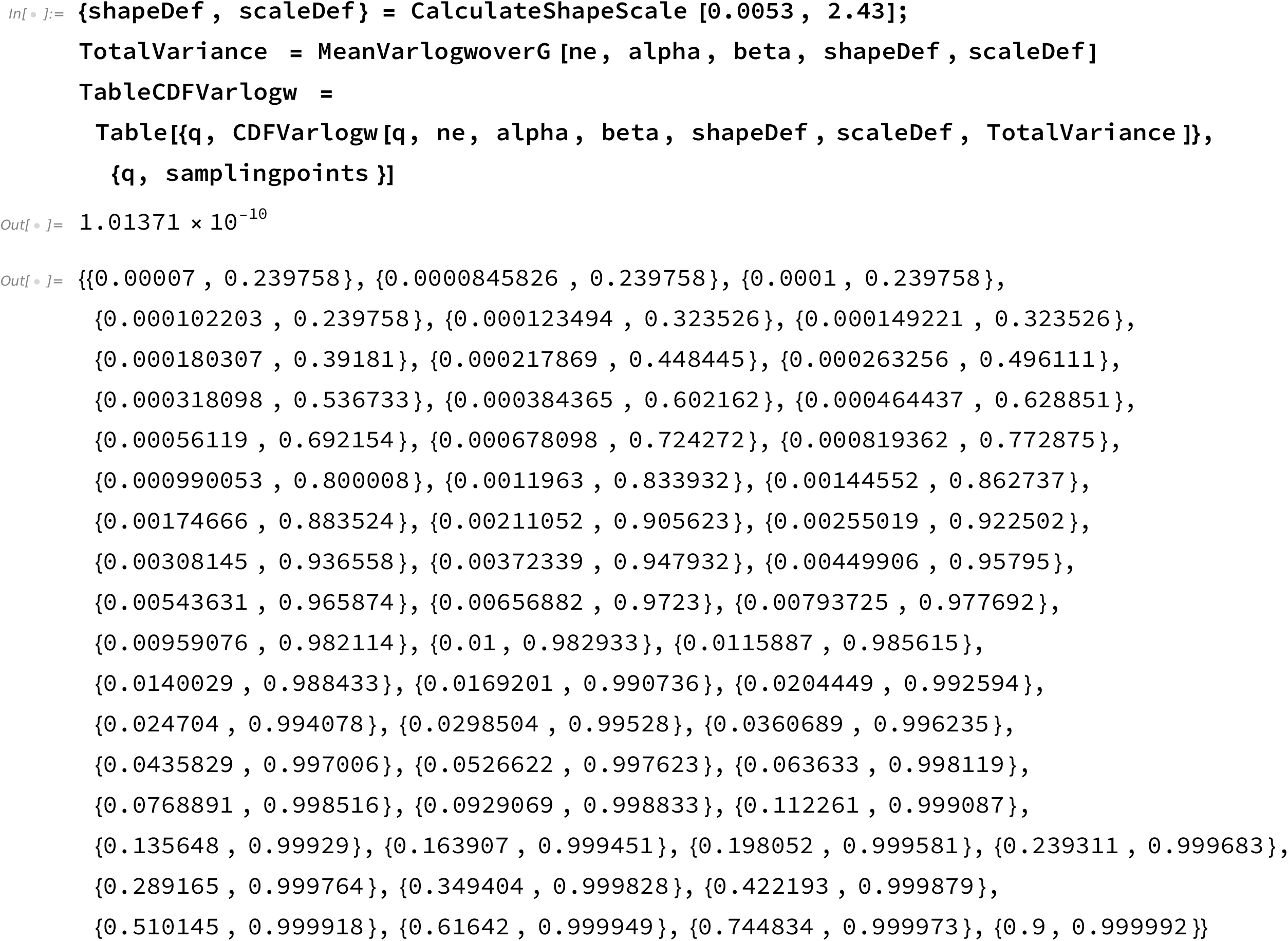
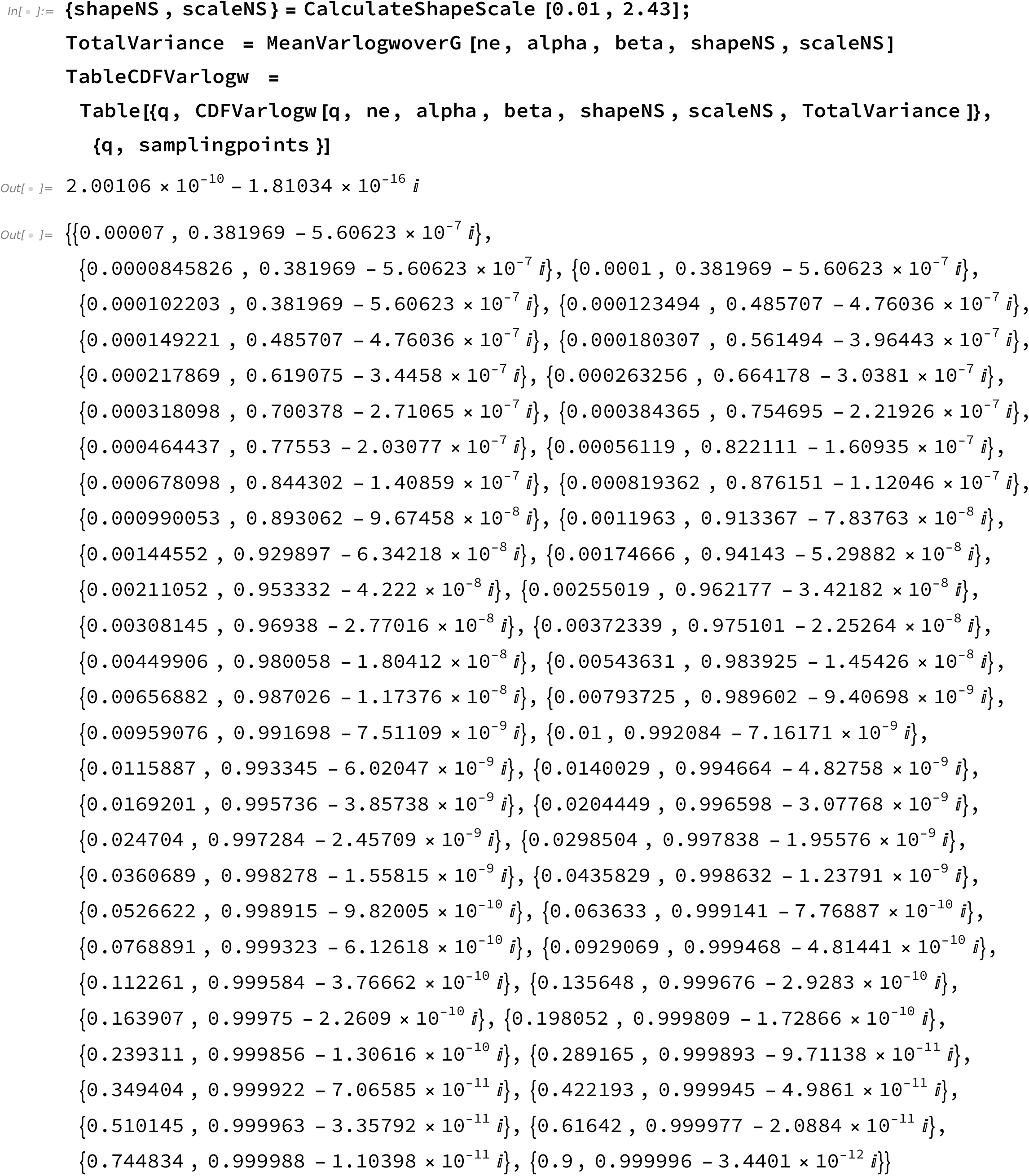
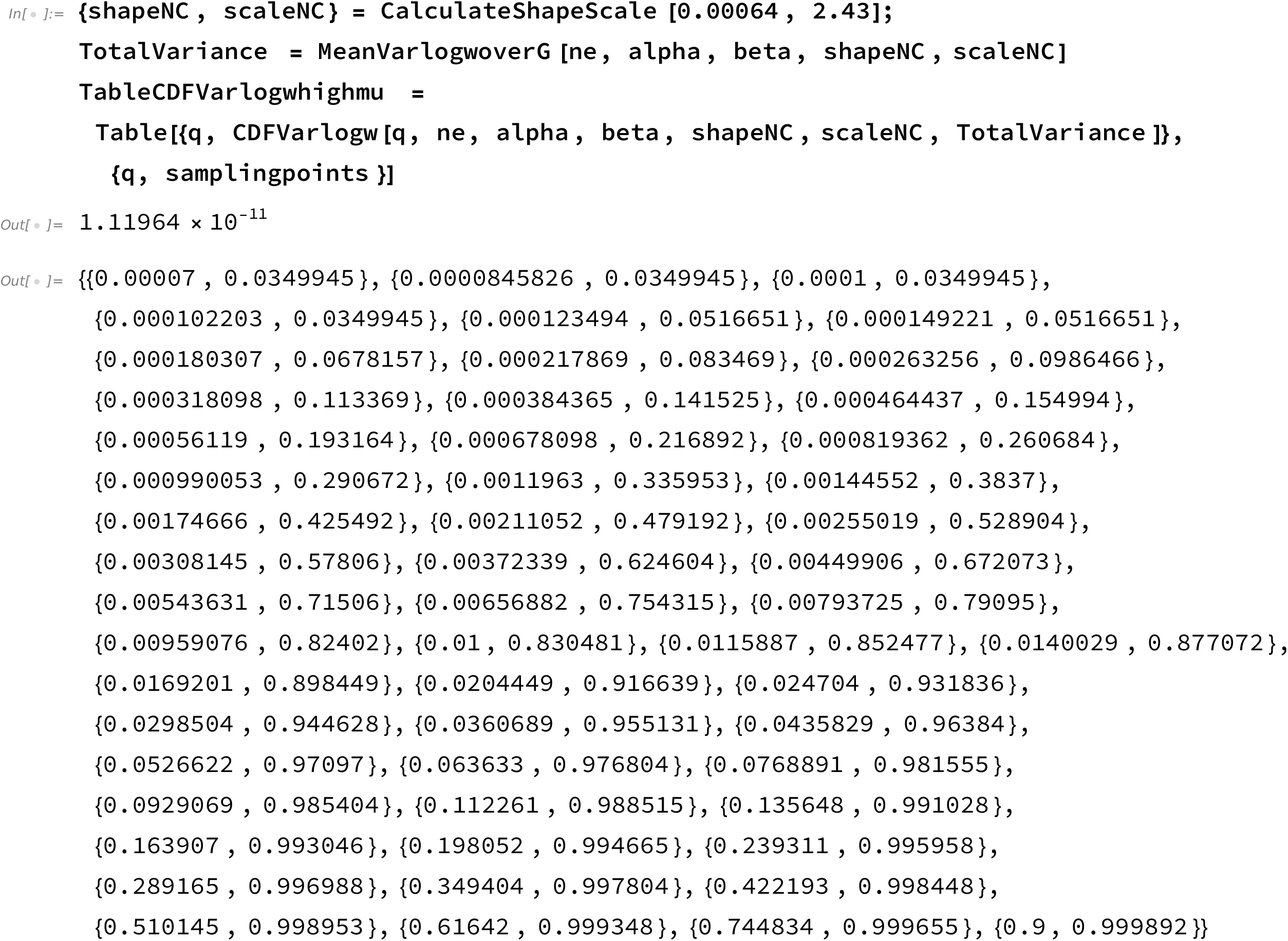

**Code figure 2 (2)**

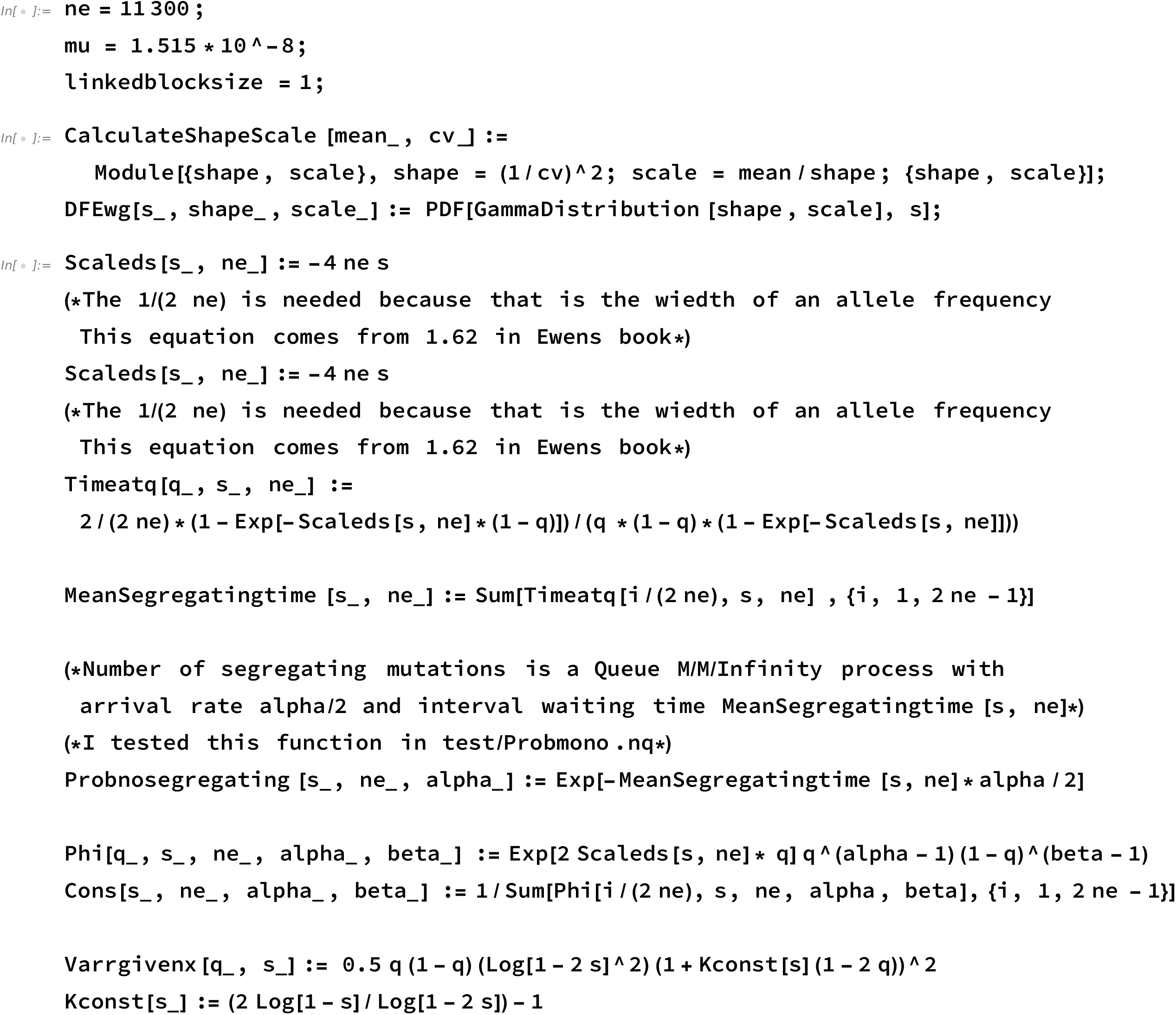
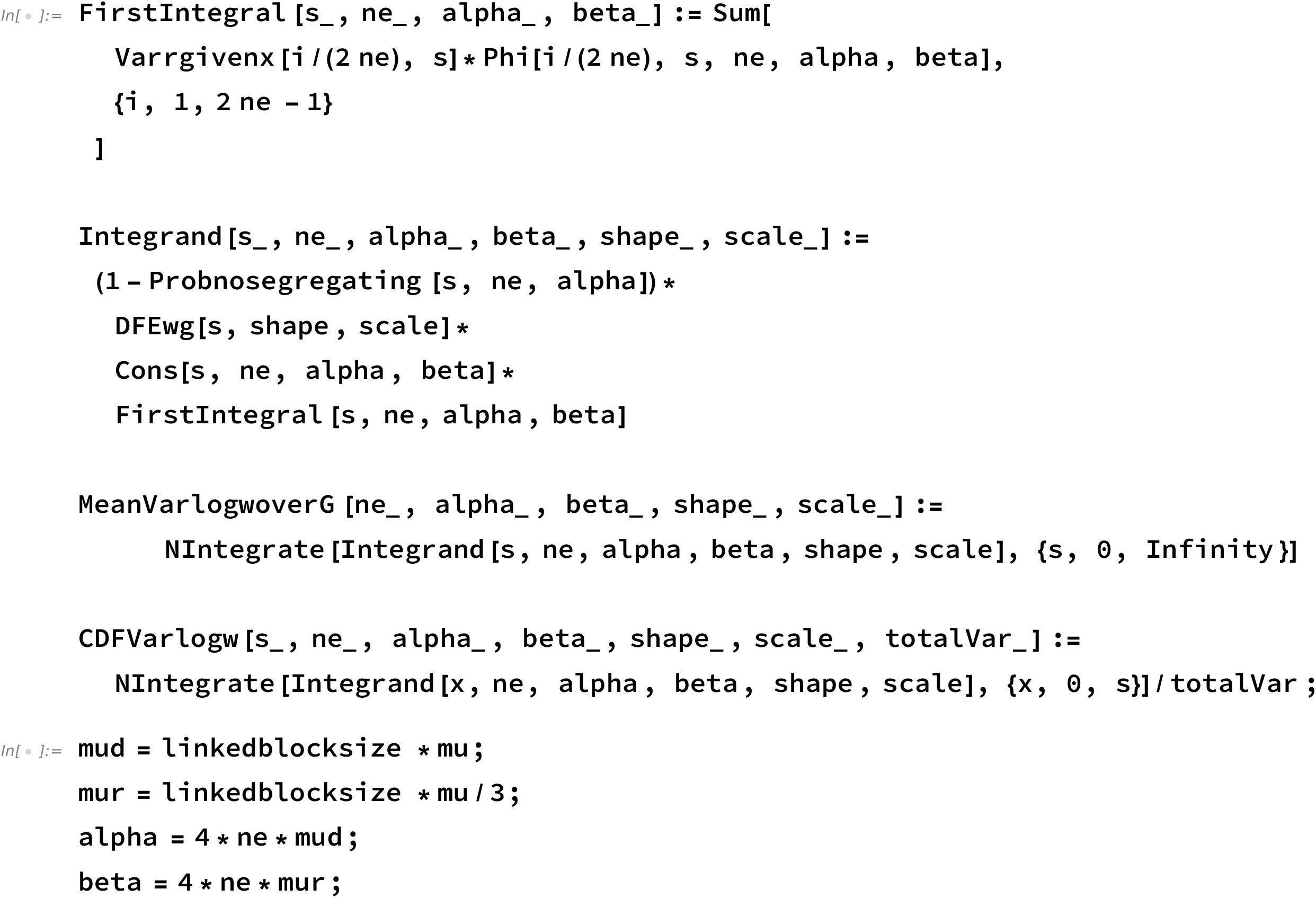
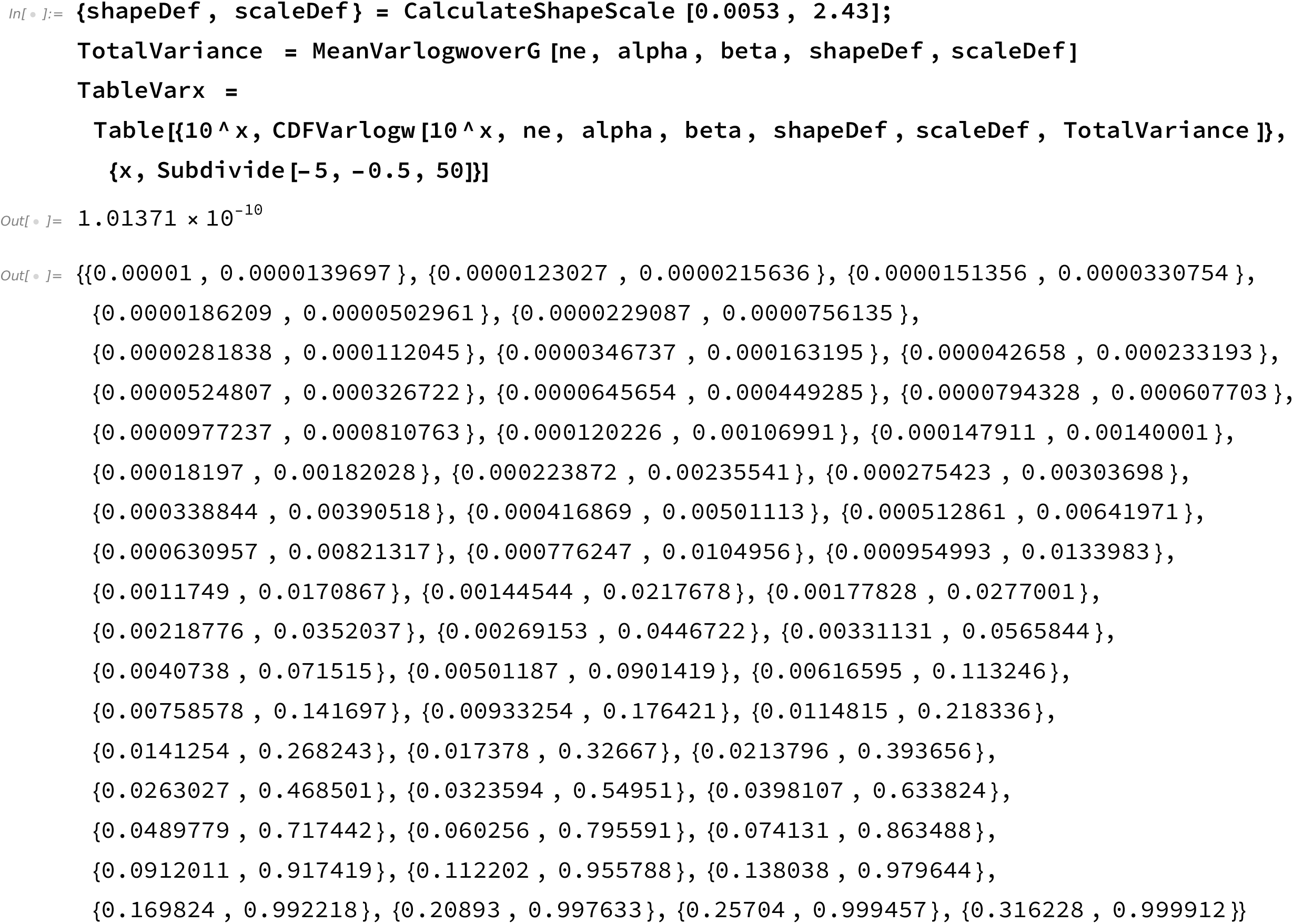
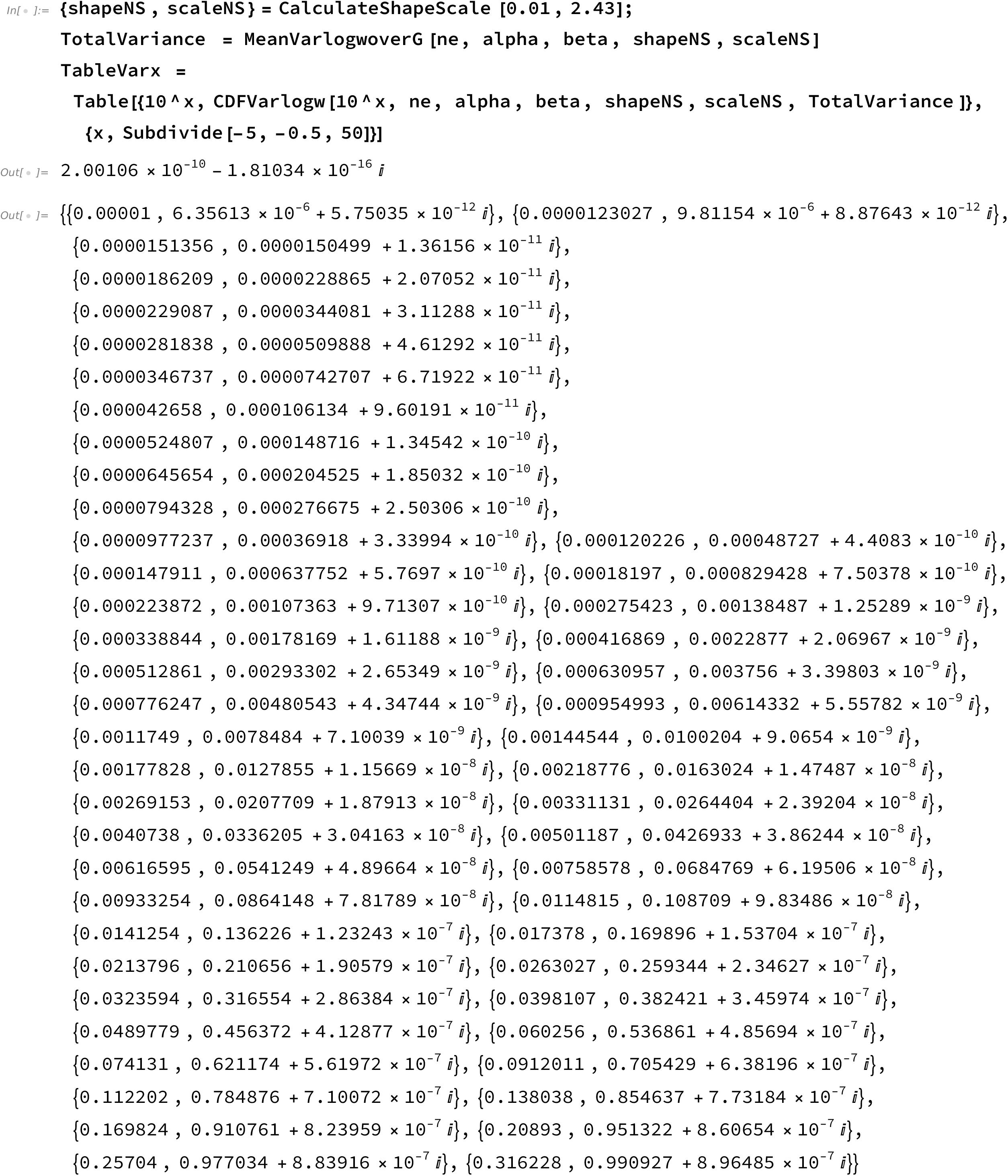
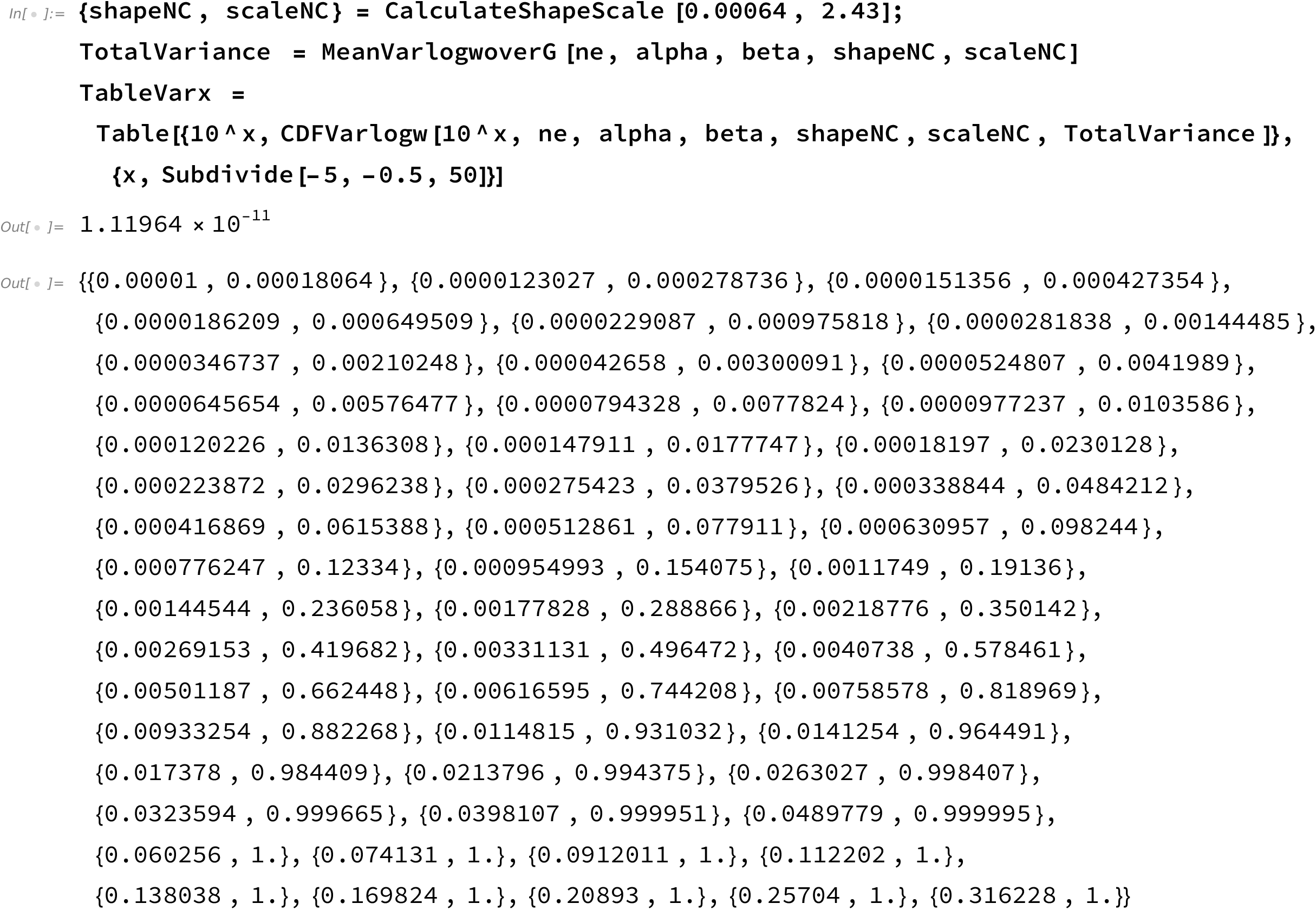

**Code figure 3**

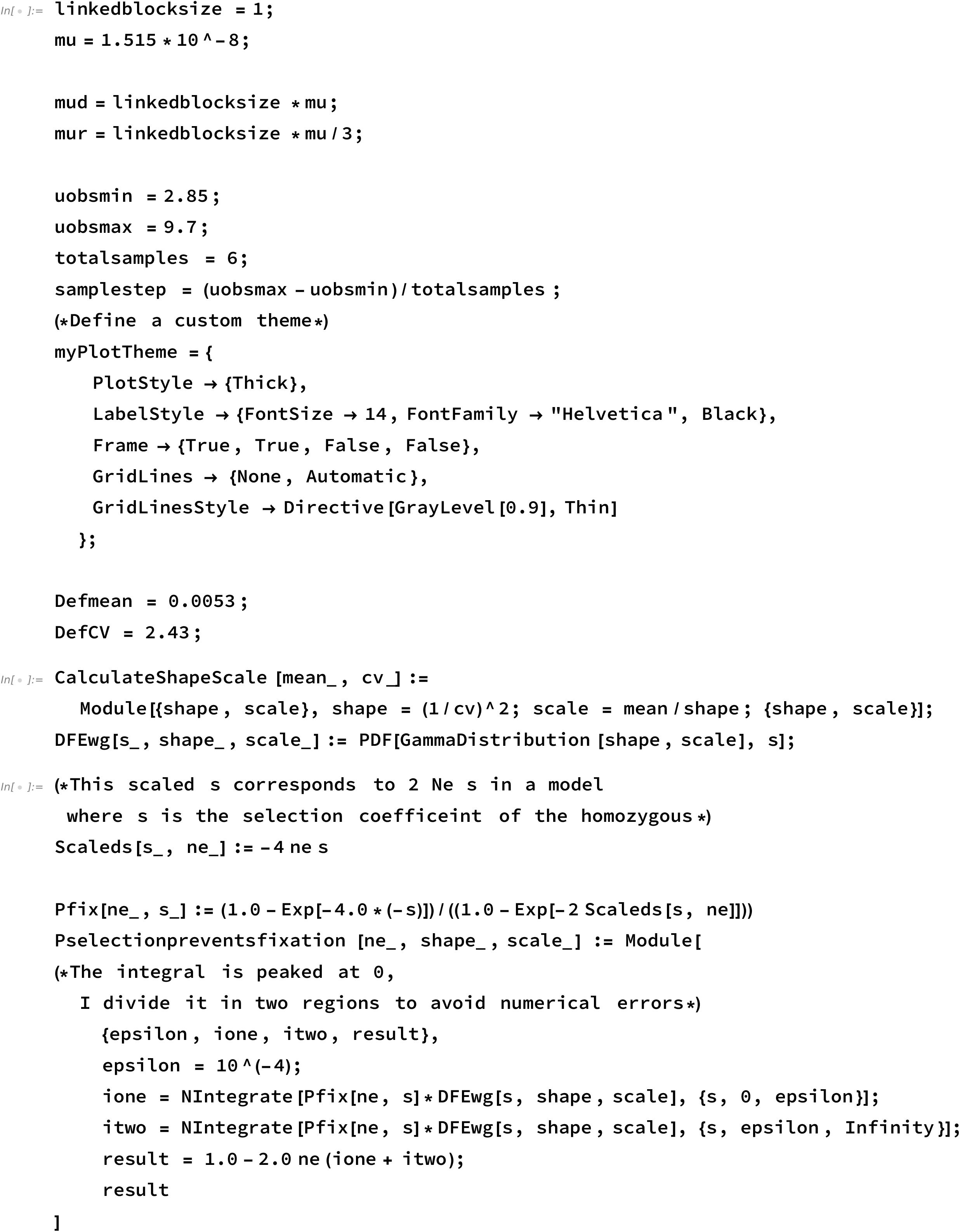
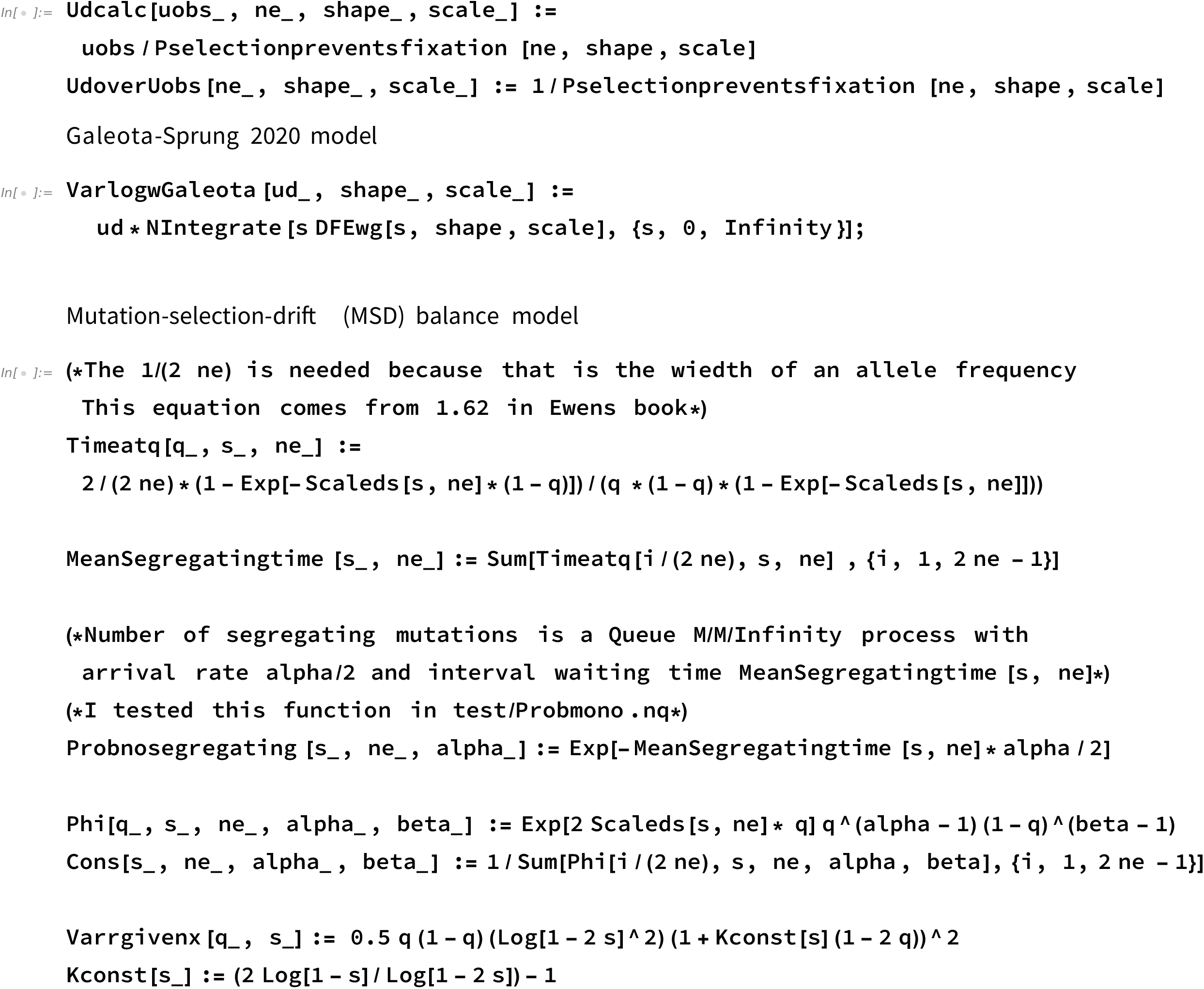
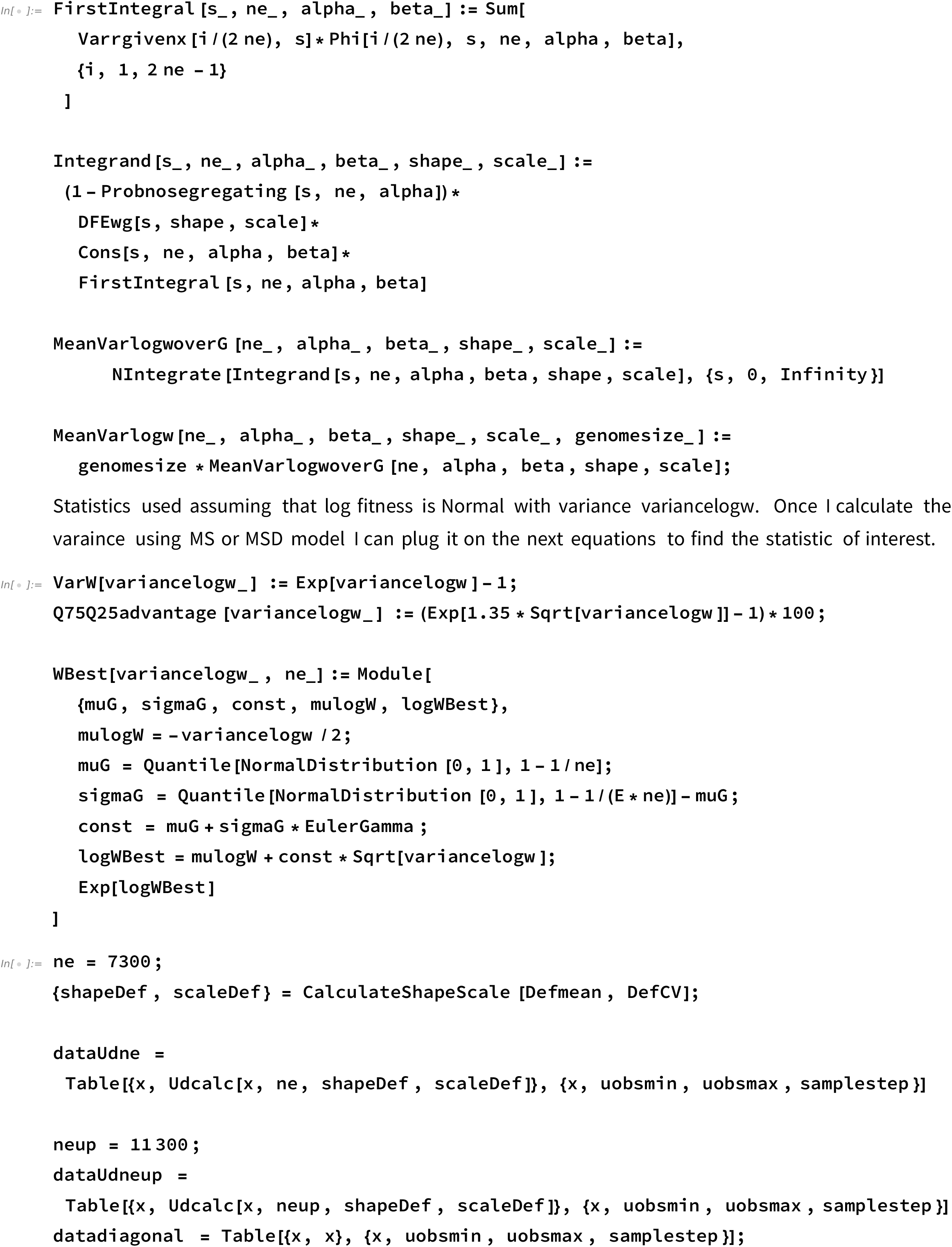
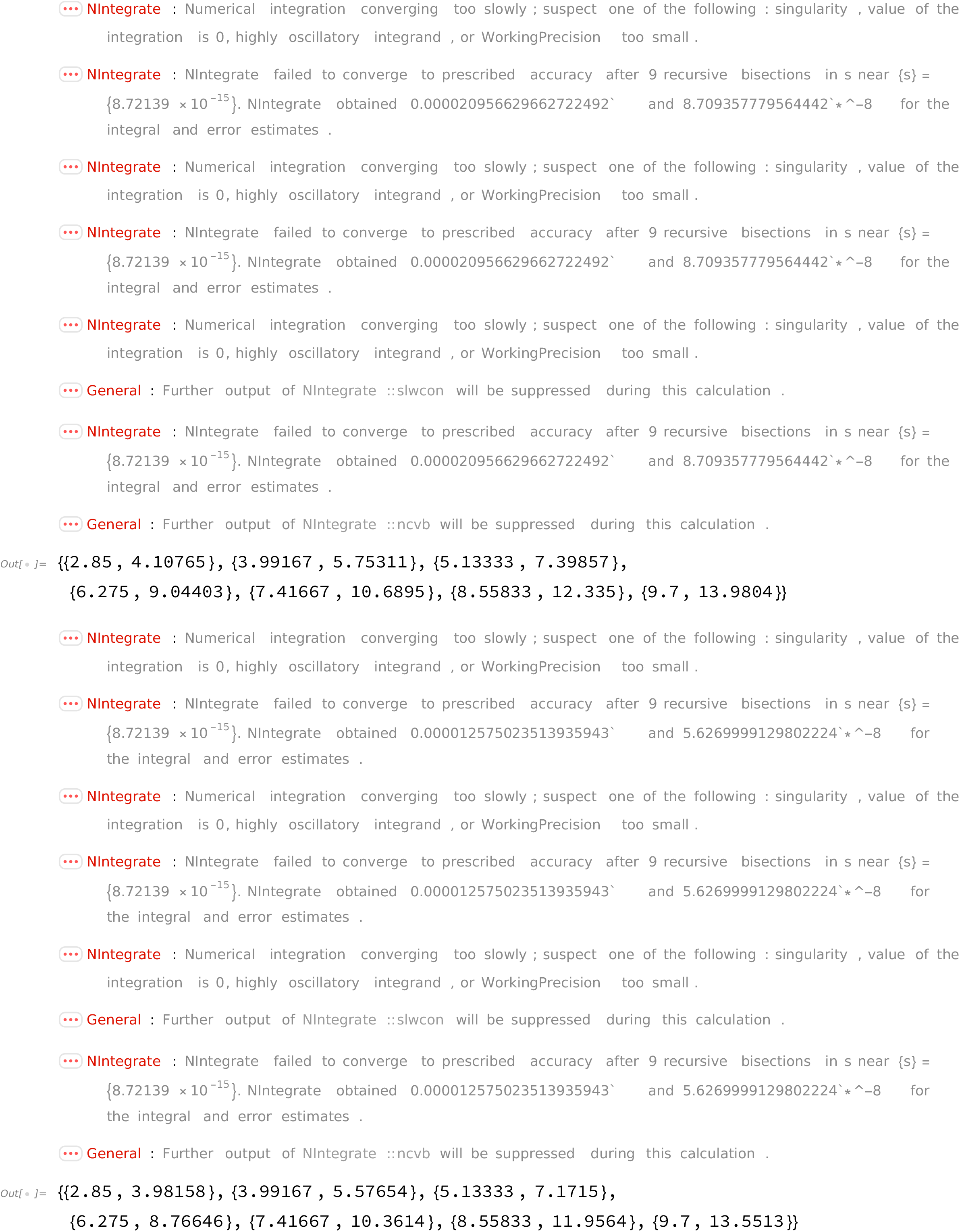
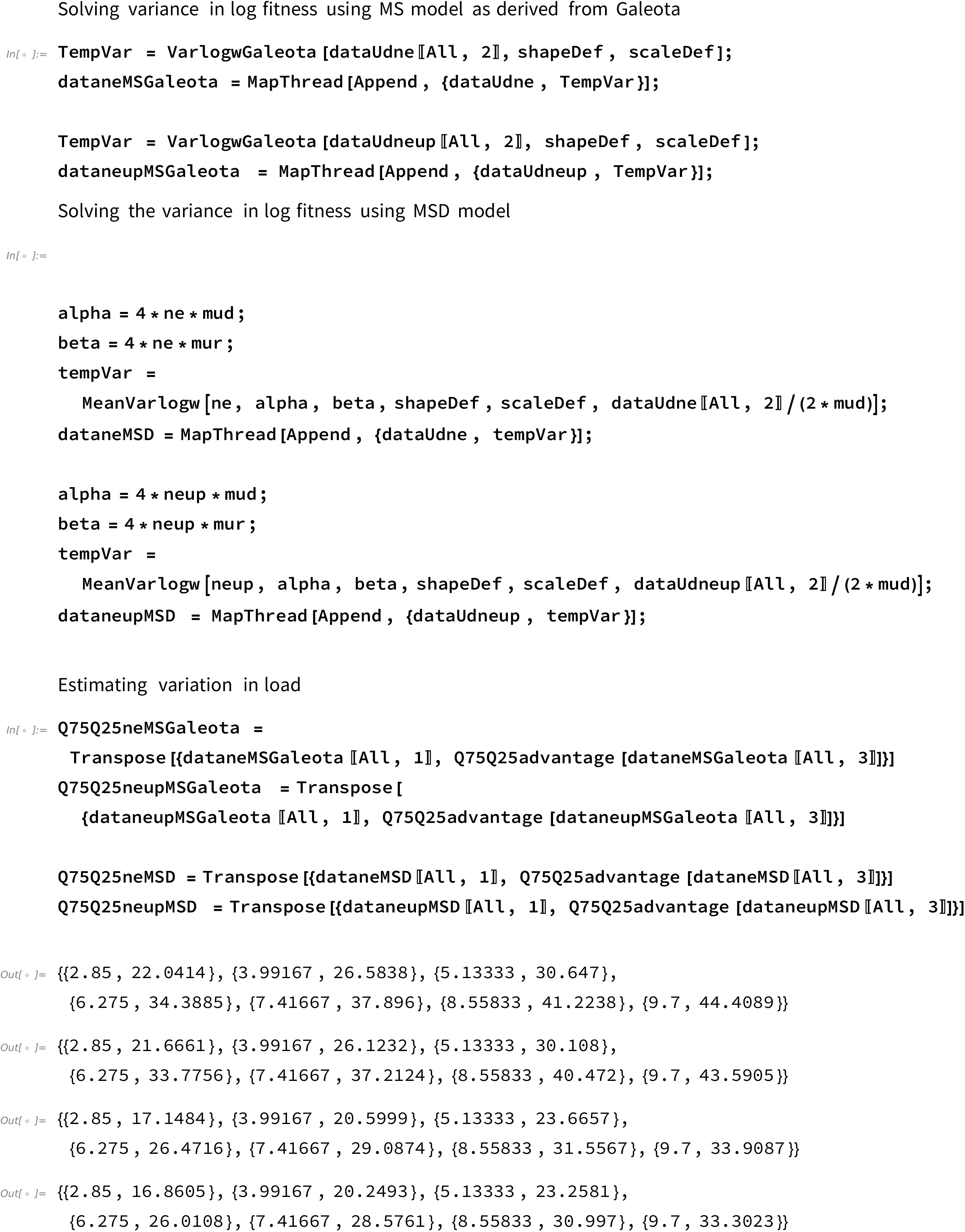
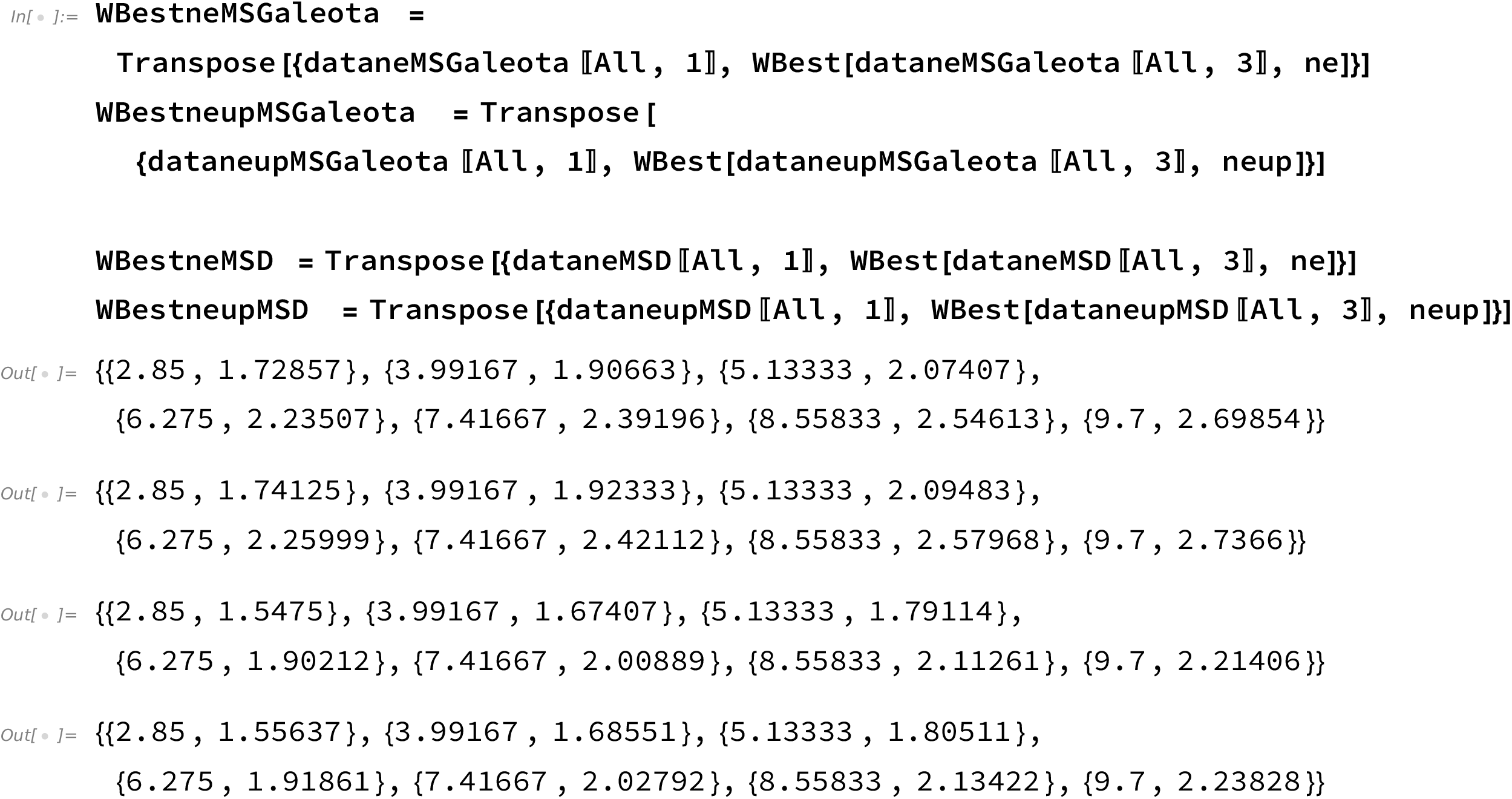

**Code figure 4**

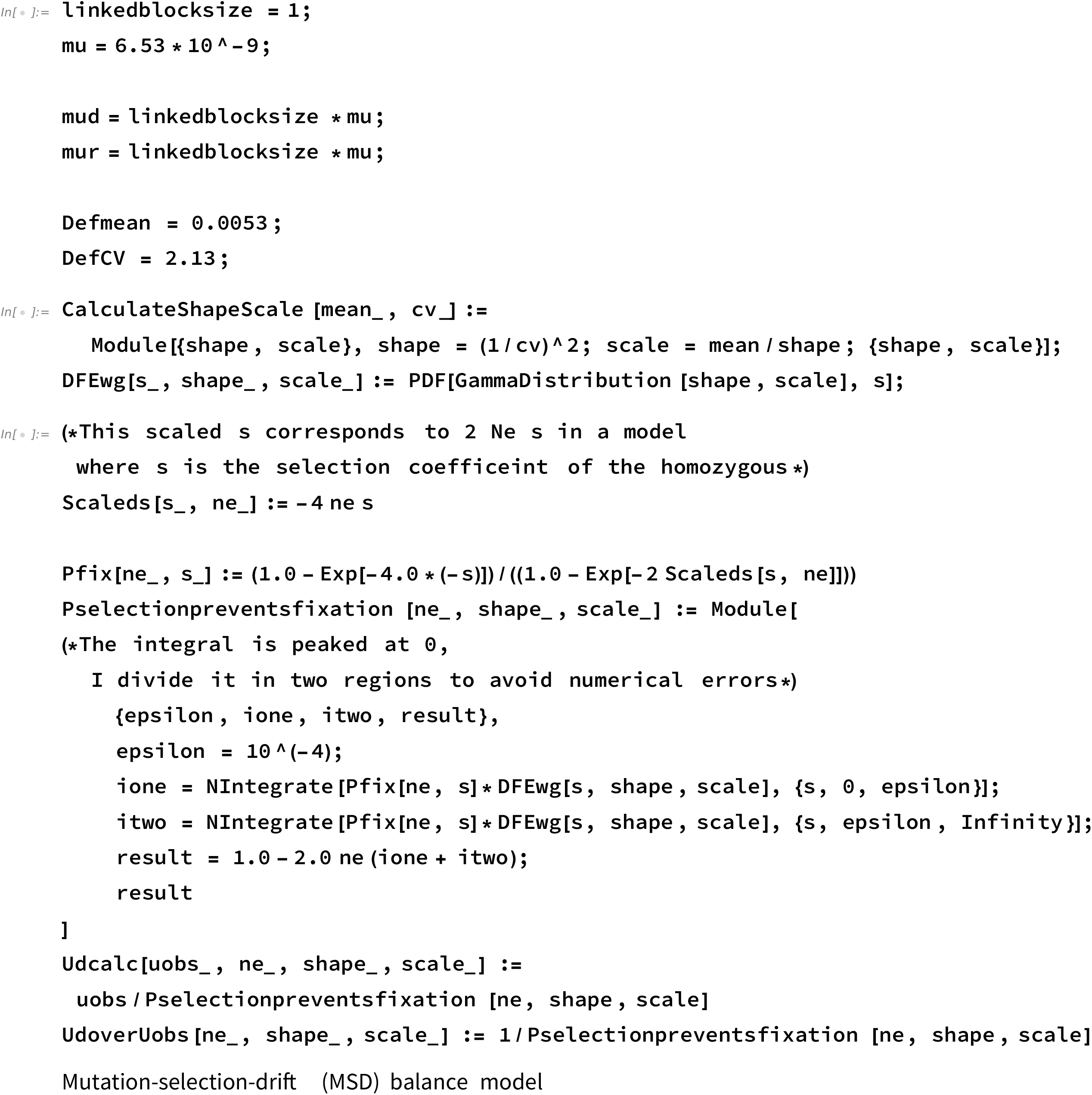
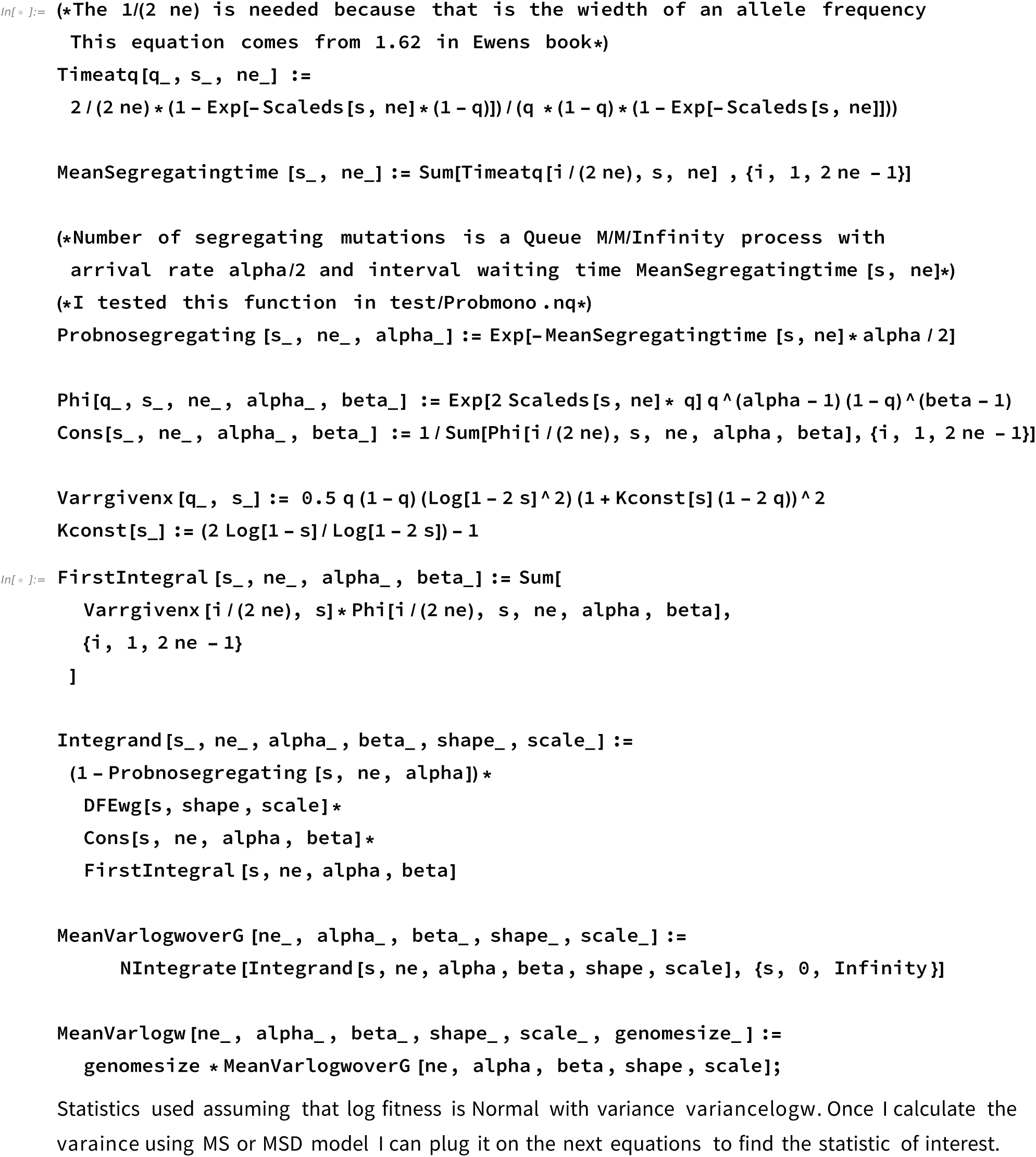
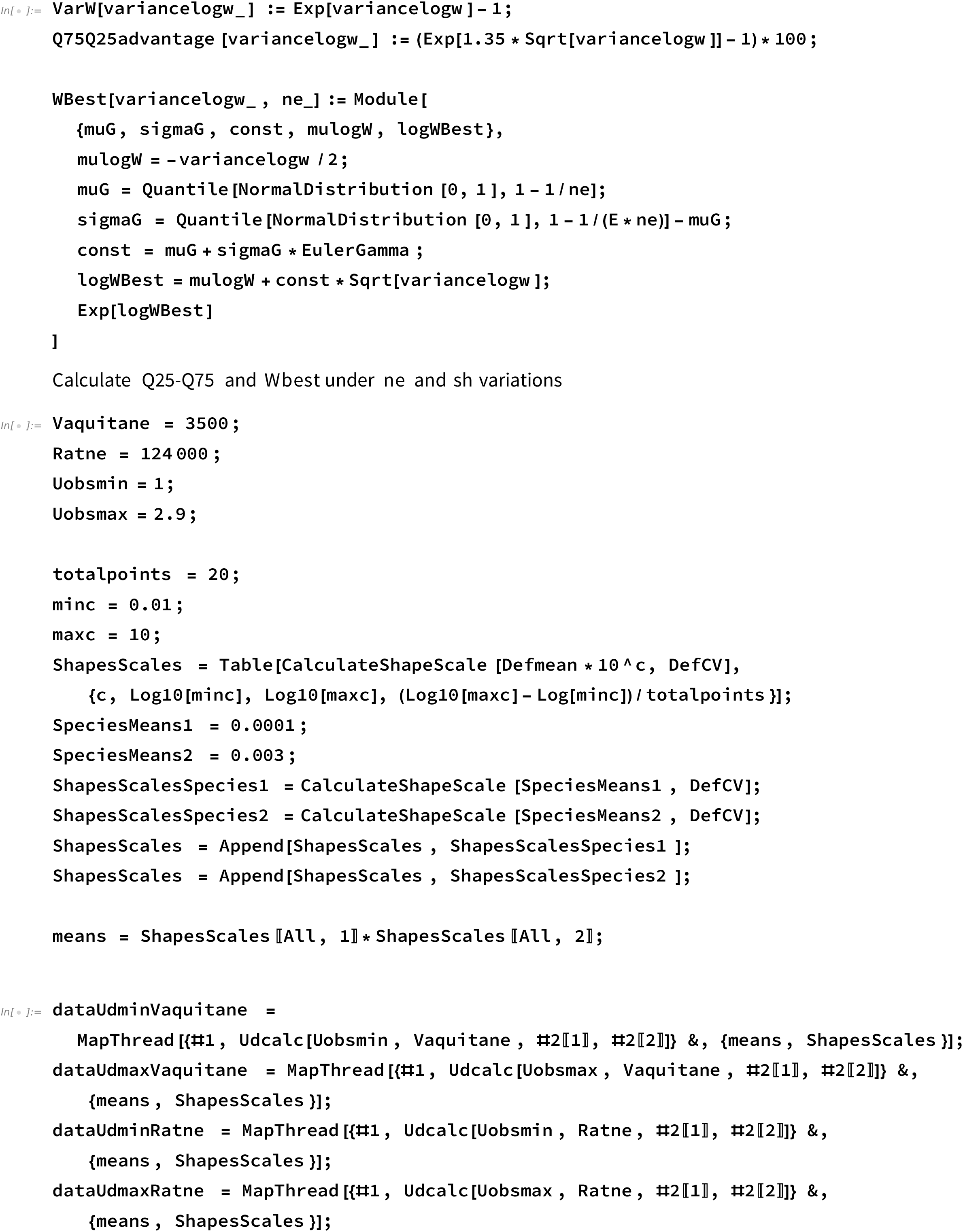
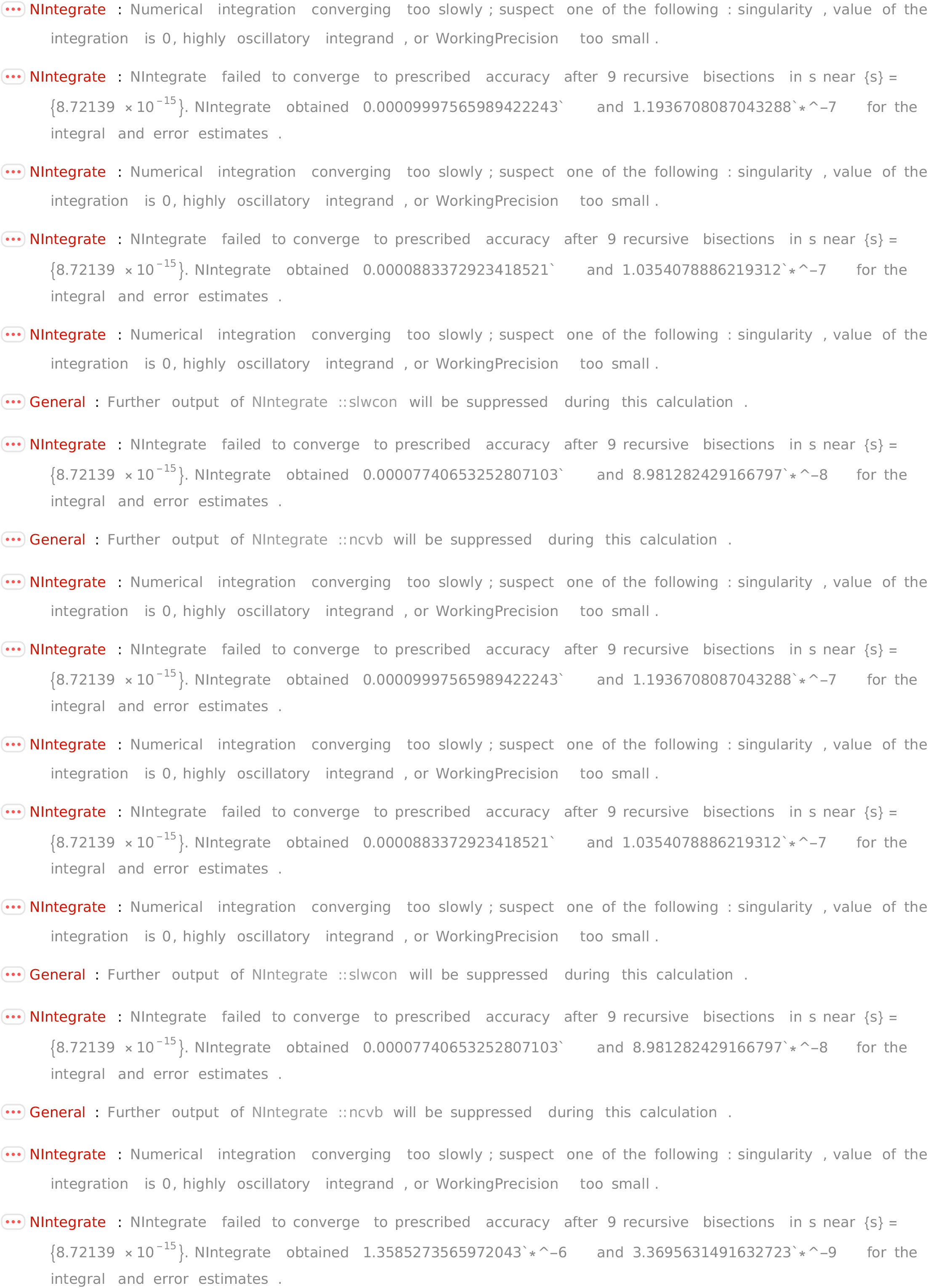
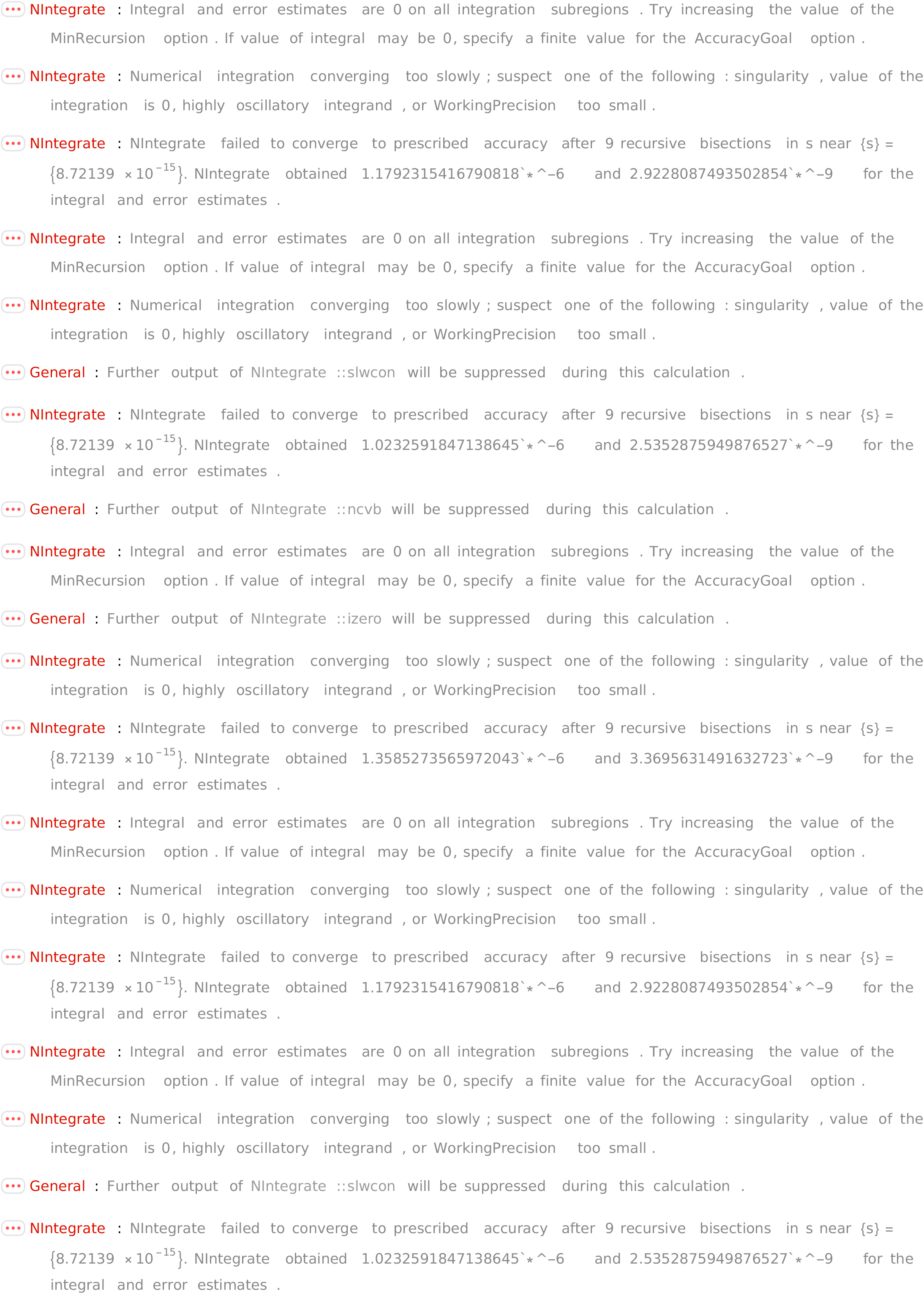
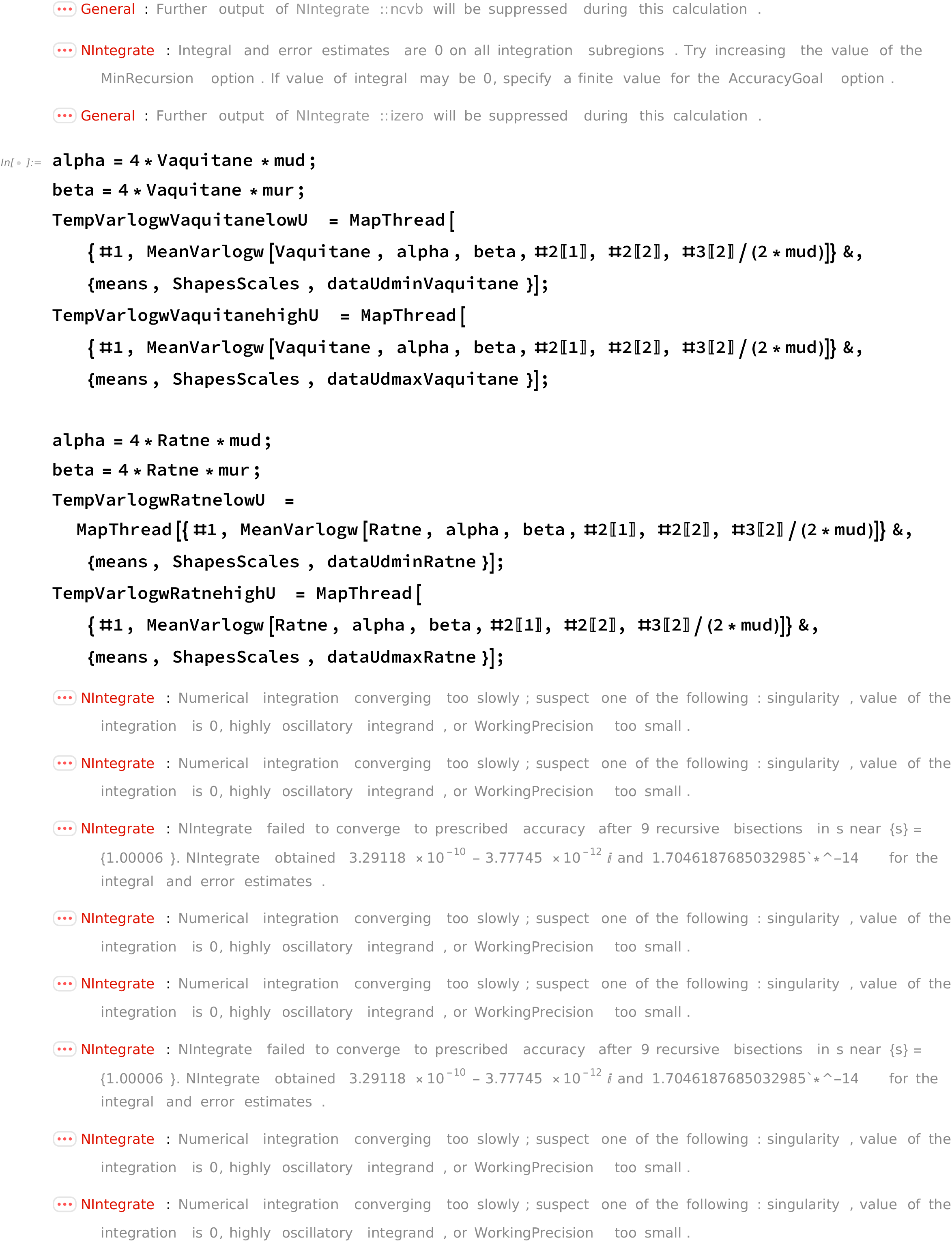
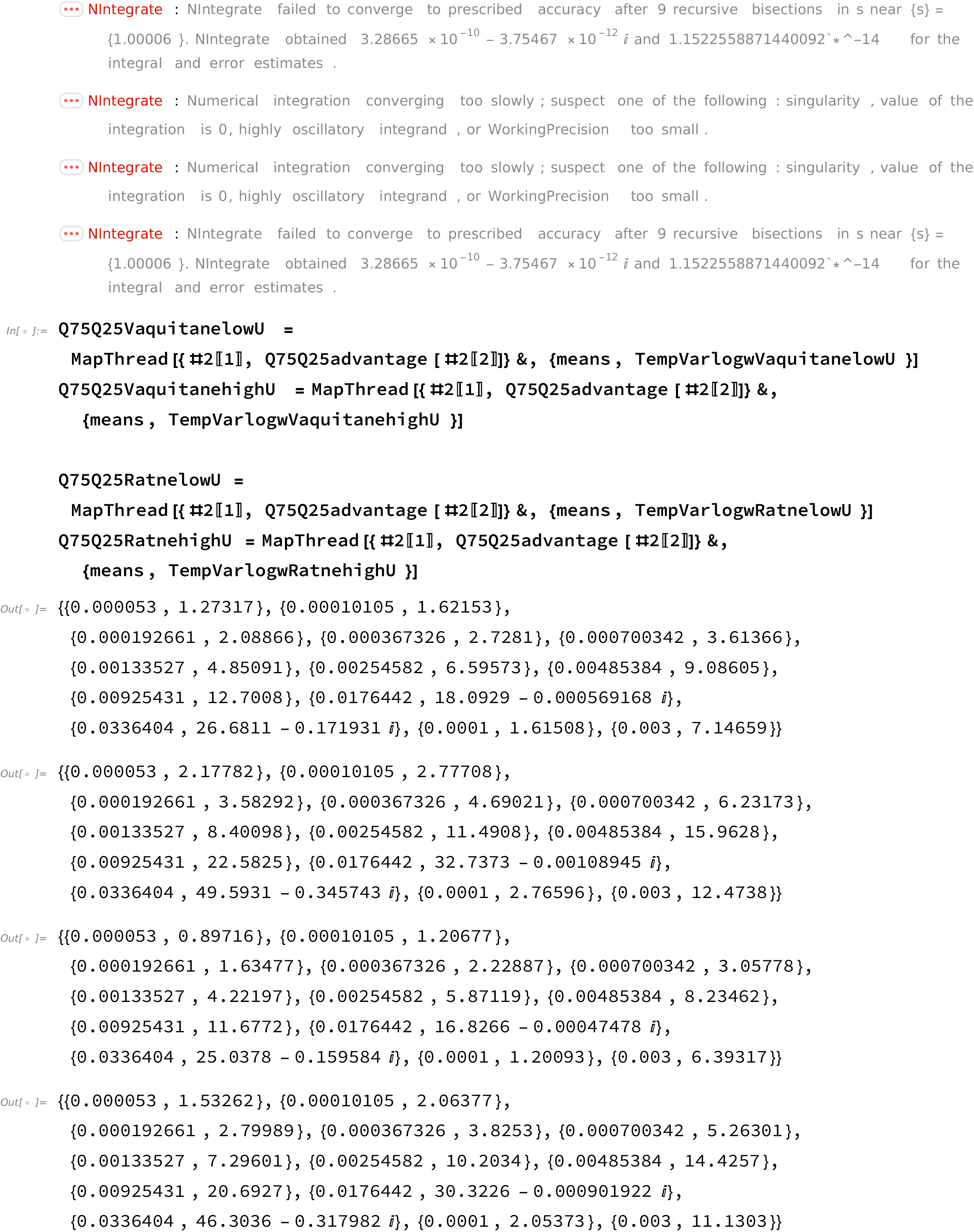
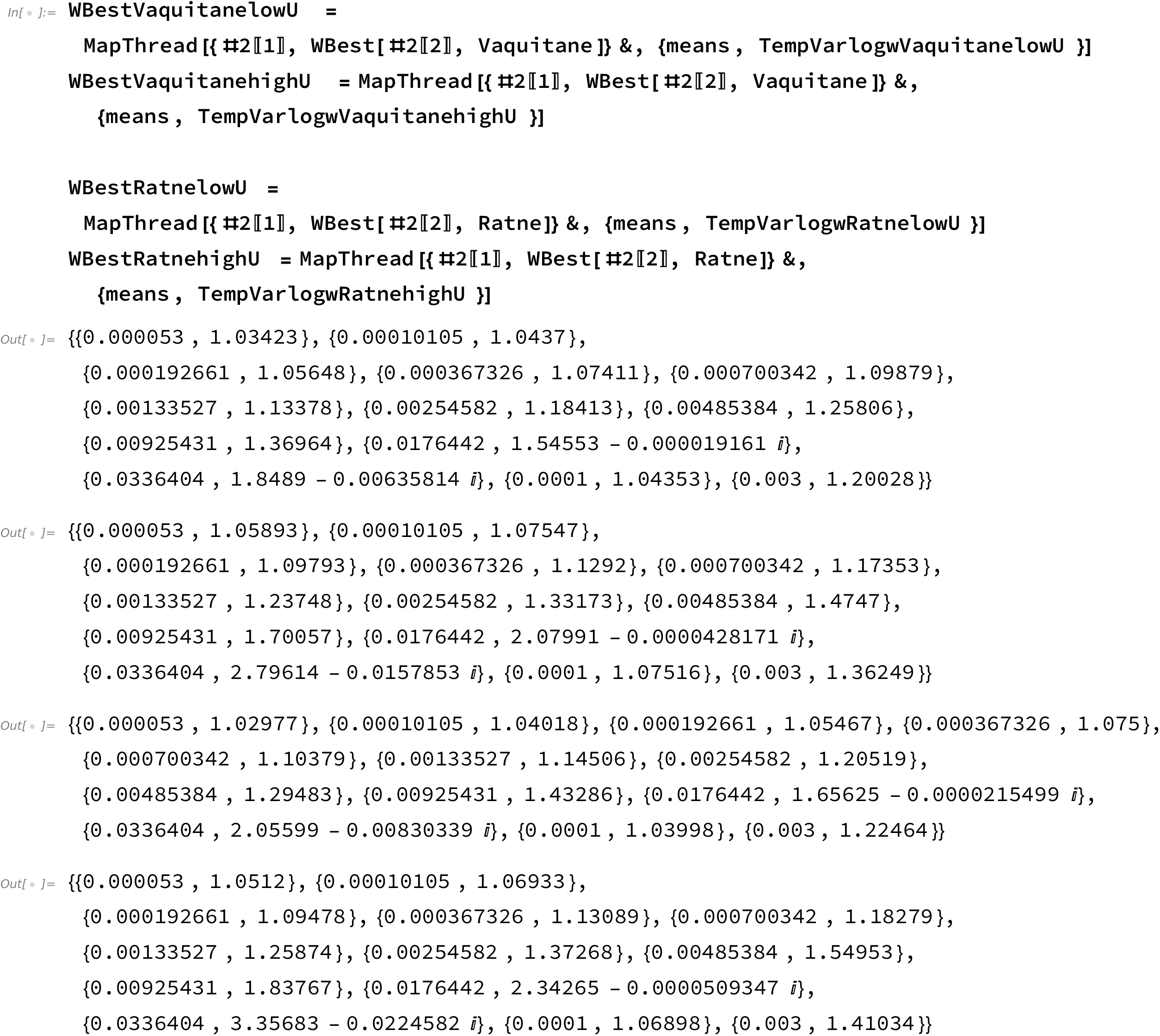

